# The interplay between homeostatic synaptic scaling and homeostatic structural plasticity maintains the robust firing rate of neural networks

**DOI:** 10.1101/2023.03.09.531681

**Authors:** Han Lu, Sandra Diaz, Maximilian Lenz, Andreas Vlachos

## Abstract

Critical network states and neural plasticity are essential for flexible behavior in dynamic environments, allowing for efficient information processing and experience-dependent learning. Synaptic-weight-based Hebbian plasticity and homeostatic synaptic scaling were considered the key mechanisms in enabling memory while stabilizing network dynamics. However, the role of structural plasticity as a homeostatic mechanism is less consistently reported, especially under activity inhibition, leading to an incomplete understanding of its functional impact. In this study, we combined live-cell microscopy of eGPF-labeled neurons in organotypic entorhinal-hippocampal tissue cultures with computational modeling to investigate the response of spine-number-based structural plasticity to activity perturbations and its interaction with homeostatic synaptic scaling. Tracking individual dendritic segments, we demonstrated that inhibiting excitatory neurotransmission does not monotonically regulate dendritic spine density. Specifically, inhibition of AMPA receptors with a low concentration of 2,3-dioxo-6-nitro-7-sulfamoyl-benzo[f]quinoxaline (NBQX, 200 nM) significantly increased spine density, while complete AMPA receptors blockade with 50 *µ*M NBQX reduced spine density. Motivated by these findings, we developed network simulations incorporating a bi-phasic structural plasticity rule governing activity-dependent synapse formation. We showed that this biphasic rule maintained neural activity homeostasis under stimulation and permitted either synapse formation or synapse loss, depending on the degree of activity deprivation. Homeostatic synaptic scaling affected the recurrent connectivity, modulated the network activity, and influenced the outcome of structural plasticity. Specifically, it reduced stimulation-triggered synapse loss by downscaling synaptic weights and rescued silencing-induced synapse loss by upscaling recurrent inputs, thus reactivating silent neurons. Our interaction between these mechanisms offers an explanation for divergent findings in the existing literature. In summary, calcium-based synaptic scaling and homeostatic structural plasticity rules compete and compensate for one another, ensuring efficient and robust control of firing rate home-ostasis.

**Significance Statement:**

- This work combined systematic computer simulations and *in vitro* experiments to explore the interplay between homeostatic structural plasticity and synaptic scaling under conditions of activity deprivation.
- We identified a non-monotonic relationship between neural activity and spine numbers, where partial inhibition of synaptic transmissions increased spine density, while complete inhibition reduced it.
- Partial inhibition led to increased spine sizes across all initial spine sizes, whereas complete inhibition selectively increased the size of relatively large spines.
- A biphasic, spine-number-based homeostatic structural plasticity (HSP) rule reconciled previously divergent experimental findings regarding activity-dependent changes in spine density.
- Using an engineering and complex systems framework, we proposed that the biphasic HSP rule incorporates a negative feedback mechanism and acts as a redundant and heterogeneous mechanism alongside the synaptic-weight-based homeostatic synaptic scaling (HSS) rule.
- By comparing simulation and experimental results, we demonstrated the necessity of HSP-HSS interplay in maintaining firing rate homeostasis.
- Both plasticity rules are driven by intracellular calcium concentration, which reflects cumulative neural activity. Thus, we propose that integral feedback control is critical in firing rate homeostasis.

## Introduction

To survive in dynamic environments, animals must quickly respond to familiar cues, such as those signaling food or predators, while remaining alert to novel stimuli. The former reflects experience-dependent learning, where subtle memory cues can trigger strong responses in the corresponding pathway during recall.^1–4^ The latter requires brain networks to maintain activity at a critical state, allowing information to be transmitted as action potentials and conveyed as neural avalanches.^5–10^ These processes represent two seemingly opposing but well-coordinated mechanisms: associative learning and firing rate homeostasis. The idea of firing rate homeostasis has been hovering in theories since the discovery of long-term synaptic potentiation (LTP),.^11^ LTP is a positive-feedback mechanism that adjusts synaptic strength among excitatory neurons (“neurons that fire together, wire together”^12^), as postulated by Donald Hebb.^13^ While its associative properties allow synapses to preserve memory traces as increased synaptic weights, it also poses the risks of driving network dynamics toward overexcitation or silence^14^ (Fig. 1Ai,Aii,Bi). *In vivo* studies in rodents have demonstrated firing rate homeostasis in the visual cortex and hippocampus, where neural activity is restored at both individual neuron^15–17^ or population level^18, 19^ within days of pertur-bation. This ability to restore network dynamics without erasing memories highlights the robustness of the complex system—the human brain network.^20–25^

**Fig. 1:**
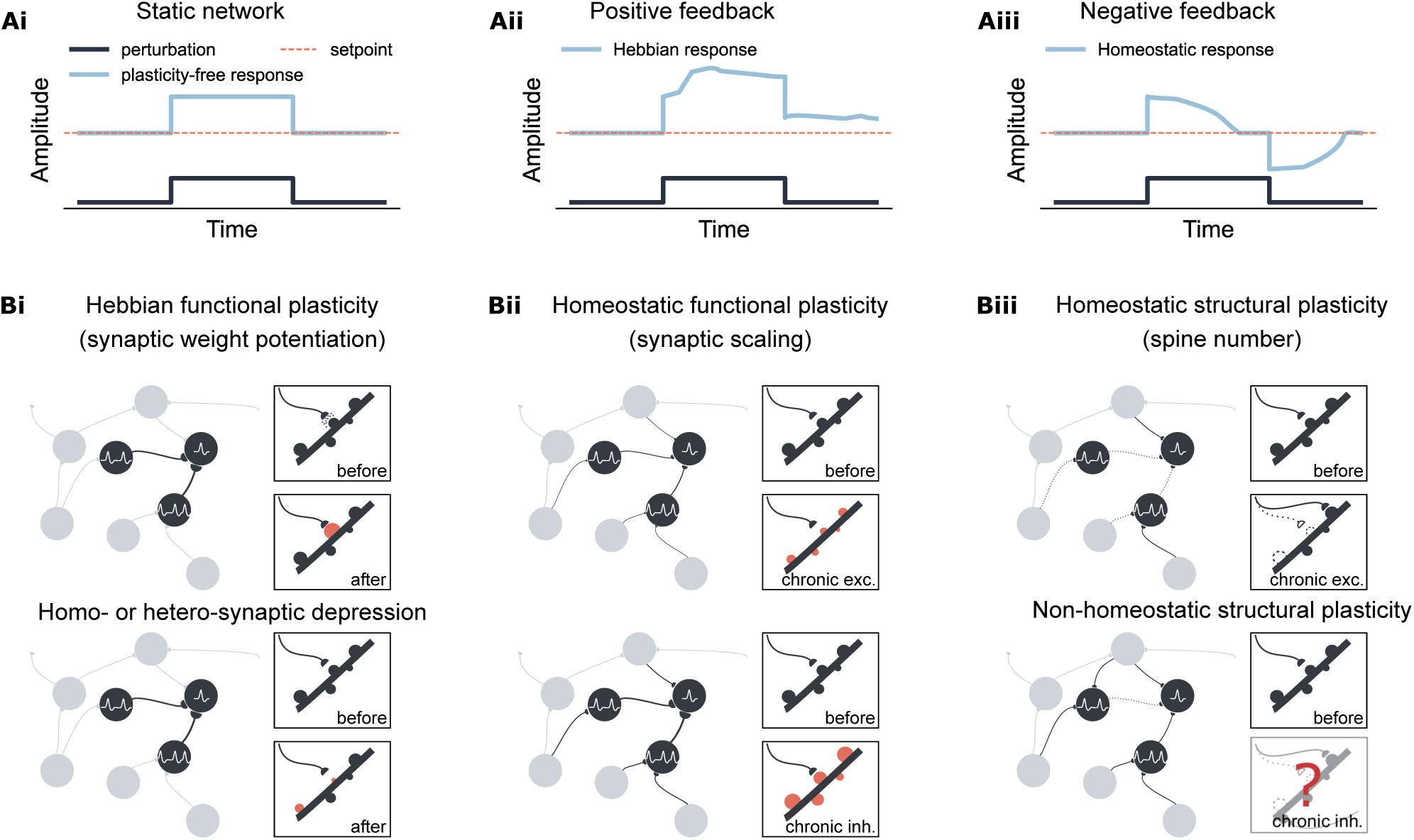
Overview of Hebbian and homeostatic plasticity. (Ai) Neural network activity is driven by external inputs in a static network. (Aii) Hebbian plasticity amplifies network responses to external inputs through a positive feedback mechanism. (Aiii) Homeostatic plasticity restores activity to a set-point via a negative feedback mechanism, essential for maintaining firing rate homeostasis. (Bi) Hebbian functional plasticity is synapse-specific, strengthening recurrent connectivity by potentiating the weights of specific synapses involved in neural activation. In contrast to synapse-specific Hebbian potentiation, synaptic depression can be induced in the same activated synapse (homosynaptic depression) or neighbor-ing synapses (heterosynaptic depression) using specific protocols. (Bii) Homeostatic synaptic scaling is cell-autonomous, involving proportional upscaling or downscaling of all input synaptic weights in response to chronic changes in neural activity. (B) Homeostatic structural plasticity is also cell-autonomous char-acterized by compensatory spine loss during chronic excitation of the postsynaptic neuron. Conversely, chronic inhibition induces divergent, often non-homeostatic, changes in spine density.

Robustness is a ubiquitous concept in many complex systems or engineering systems, meaning that when external perturbations forcefully change its dynamics, the system adapts to return to its original attractor or moves to a new attractor state that preserves its functions.^23, 24^ Essential features of robustness include negative feedback control, redundancy, and heterogeneity.^20–25^ For instance, homeostatic synaptic scaling exemplifies negative feedback strategy^26^ (Fig. 1Aiii,Bii), as neurons proportionally adjust their synaptic weights in a compensatory manner in response to long-term changes in neural activity,^26^ membrane potential,^27^ or intracellular calcium concentration.^28^ Redundancy and heterogeneity encompass various mechanisms that achieve the same goal, providing backup in case of failure. Although redundancy and heterogeneity have been recognized in biological systems in the form of genetic buffering and convergent molecular circuits,^23, 24, 29^ their representation in activity-dependent plasticity has been less explicitly discussed. Induction of homosynaptic long-term depression (LTD)^30^ under the Bienenstock–Cooper–Munro (BCM) rule^31^ or spike-timing-dependent plasticity (STDP) rule,^32–35^ or induction of heterosynaptic LTD via synaptic tagging and capture^36^ contribute to stabilizing firing rates. Inhibitory plasticity also helps restore the balance between excitation and inhibition.^37–39^ These mechanisms, which modulate synaptic transmission locally or globally, exhibit redundancy in function and heterogeneity in the implementation in reference to homeostatic synaptic scaling, serving the same purpose of stabilizing the network dynamics.

However, it remains unclear whether such redundancy extends to the structural substrate of synaptic transmission. Structural plasticity encompasses changes in sizes or numbers of dendritic spines and axonal boutons, synapse numbers, and network connectivity, all of which are critical to functional transmission and memory capacity.^40^ The homeostatic structural plasticity (HSP) model, which posits that synapse formation and elimination are homeostatically regulated,,^41–45^ has expanded the potential for modelling network reorganization in response to diverse perturbations.^19, 46–49^ Nonetheless, experimental results present a more complex narrative (Fig. 1Biii): Studies often find spine loss during synaptic downscaling, while inconsistent and mostly non-homeostatic spine loss is seen in upscaling experiments under chronic activity deprivation.^50^ This complexity underscores the need for a deeper understanding of activity-dependent structural plasticity, particularly the rules governing spine number changes. Given the range of experimental conditions, including monocular deprivation, lesions, and pharmacological treatments, we hypothesized that: (i) the relationship between neural activity and spine numbers is non-monotonic, and (ii) homeostatic synaptic scaling might confound the interpretation of structural plasticity rules based on the inconsistent experimental results.

In this study, we used two concentrations of the competitive AMPA-receptor antagonists NBQX to gradually block neural activity. Using whole-cell patch-clamp recordings and time-lapse imaging, we found that activity deprivation affects dendritic spine numbers and sizes in a non-monotonic manner. A low concentration of NBQX (200 nM) increased spine numbers, while a higher concentration (50 *µ*M) reduced them. These findings guided us to establish a spiking neural network model with a biphasic, synapse-number-based HSP rule, which successfully captured both homeostatic and non-homeostatic changes in response to varying degrees of activity perturbations. Additionally, we implemented a monotonic, synaptic-weight-based homeostatic synaptic scaling (HSS) rule based on the literature.^51^ Although largely redundant to the biphasic HSP rule, the HSS rule facilitates a shift from non-homeostatic synapse loss to homeostatic synapse regeneration by adjusting recurrent connectivity in silenced networks. Our results highlight the redundancy and complementarity between these two rules, underlining the combined importance of HSP and HSS in enabling robust and efficient adaptation of network activity.^25, 52, 53^

## Results

### Informing a non-monotonic structural plasticity rule with gradual activity blockade experiments

The variability in experimental results regarding structural plasticity under chronic activity deprivation^50^ could arise from either non-monotonic, activity-dependent regulation of spine numbers, concurrent changes in synaptic weights and firing rates due to synaptic scaling, or a combination of both. To comprehensively address this, we established computer simulations that integrate the synapse-number-based structural plasticity rule with the synaptic-weight-based synaptic scaling rule. These activity-dependent models function as integral feedback mechanisms linked by the same intracellular calcium concentration *C*(*t*) and setpoint value *ɛ* (Fig. 2A), which monitors neural activity and influences synaptic weights or synapse numbers. This approach eliminated the need for manual time-scale alignment. Structural plas-ticity models vary, with some suggesting neural activity affects neurite growth and retraction, thereby influencing synapse formation and loss either monotonically^19, 46, 47^ or non-monotonically.^41, 43–45^ Given the need for additional experimental data to inform model selection, we conducted experiments to test whether spine density responds monotonically to changes in neural activity.

**Fig. 2:**
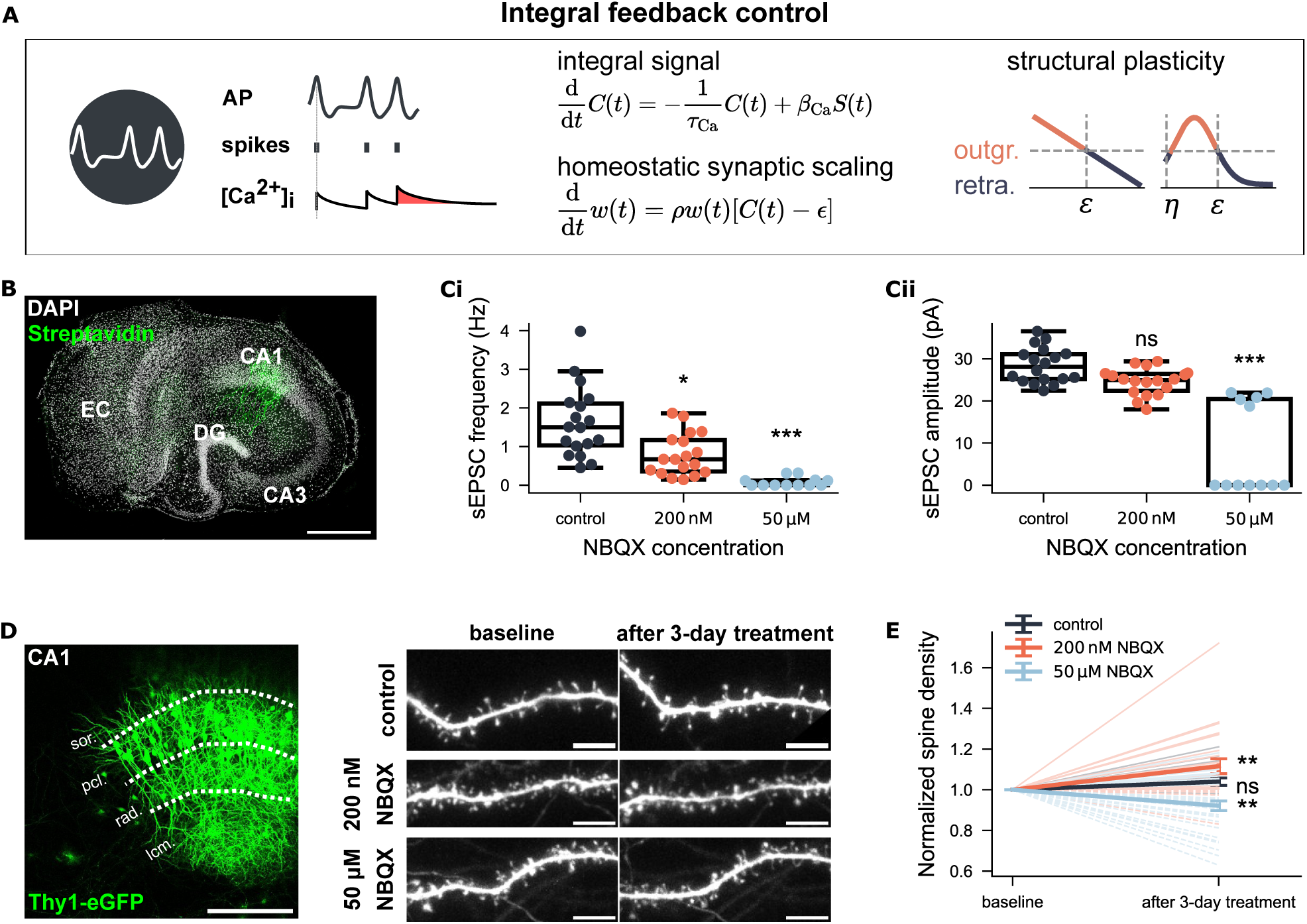
Non-monotonical modification of spine numbers following three-day synaptic inhibi-tion via NBQX informs a biphasic structural plasticity rule. (A) The integral feedback control framework for homeostatic synaptic scaling and structural plasticity. Both rules use intracellular calcium concentration ([Ca^2+^]_i_, *C*(*t*)) to track neural activity (AP, action potential, *S*(*t*)). Calcium concentra-tion is updated each time via calcium influx (*β*_Ca_) upon the emission of a post-synaptic action potential and decays with a time constant *τ*_Ca_. Synaptic scaling follows a weight-dependent rule, multiplicatively adjusting the synaptic weight *w*(*t*) with a scaling factor *ρ*, based on the discrepancy from the setpoint *ɛ* of calcium concentration. Structural plasticity uses the same guidance signal (i.e., calcium concentration) to regulate the growth and retraction of axonal boutons and dendritic spines, aiming to reach the same setpoint value. Two versions of the structural plasticity rule are shown: one with a linear dependency and the other with a non-linear dependency on intracellular calcium concentration. (B) Example CA1 pyramidal neuron recorded from entorhinal-hippocampal tissue culture to assess the effects of different NBQX concentrations. Scale bar: 500 *µ*m. (Ci-Cii) Group data of spontaneous excitatory postsynaptic currents (sEPSC) in three groups (*N* = 18 for the control group, *N* = 18 for the 200 nM NBQX-treated group, *N* = 12 for the 50 *µ*M NBQX-treated group). (D) Example Thy1-eGPF culture and dendritic segments from the stratum radiatum (rad.) before and after three-day treatment. Scale bars: 200 *µ*m and 5 *µ*m. (E) Spine density at baseline and after the three-day treatment. All values are normalized to the corresponding baseline values. Light-shaded lines show raw data (solid lines represent increased spine density; dashed lines represent decreased density). Dark-shaded lines with error bars represent the means and standard error of the means (s.e.m.s) for each group. (*N* = 19 for the control group, *N* = 24 for the 200 nM-treated group, *N* = 33 for the 50 *µ*M-treated group.)

We used two concentrations of the AMPA receptor antagonist NBQX – 200 nM and 50 *µ*M – and studied CA1 pyramidal neurons in organotypic entorhinal-hippocampal tissue cultures (Fig. 2B). Whole-cell patch-clamp recording of AMPA receptor-mediated spontaneous excitatory postsynaptic currents (sEPSCs) revealed a significant reduction in mean sEPSC amplitude with 200 nM NBQX and a near-to-complete blockade of excitatory synaptic transmission with 50 *µ*M NBQX (Fig. 2Ci-Cii). Both NBQX concentrations were applied to Thy1-eGFP cultures for three days to induce chronic activity inhibition. Individual dendritic segments (side branches of apical dendrites in the stratum radiatum) were imaged before and after treatment using time-lapse microscopy (Fig. 2D). The analysis showed that 200 nM NBQX increased spine density (*p* = 0.003, Wilcoxon test), while 50 *µ*M NBQX reduced spine density (*p* = 0.008, Wilcoxon test), compared to their baseline densities (Fig. 2E). No significant changes in spine density were observed in the control cultures (*p* = 0.06, Wilcoxon test). These results demonstrated a non-monotonic relationship between neural activity and spine numbers under activity deprivation: partial inhibition increased spine numbers, while complete inhibition led to a reduction.

### Stabilizing and characterizing a biphasic structural plasticity rule

Since changes in spine numbers or density are functionally linked to synapse formation, loss, and rewiring, we aimed to establish structural plasticity rules in point neurons and study synaptic rewiring in a large, topology-free spiking neuron network, thereby bypassing the complexities of neural morphology (Fig. 3A). For comparison, we first simulated the linear growth rule (Fig. 3Bi). To capture the observed non-monotonic dependency in a simplified manner, we adopted a Gaussian-shaped growth rule for synaptic element numbers with two setpoints. By setting the first setpoint at either *η* = 0 or *η >* 0, we generated two variants of the Gaussian rule (Fig. 3Bii-Biii). We then grew three neural networks where the synapses among excitatory neurons were governed by these three rules, respectively (Fig. 3C). As reported in previous studies,^19, 45, 46, 48^ the linear-rule-guided network developed smoothly to a homogeneous sparsely connected network (10% of connection probability, equivalent to approximately 1 000 excitatory synapses per neuron) with an average firing rate near the target value of *ɛ* = 7.9 Hz (Fig. 3Di). The Gaussian rule with a zero setpoint (*η* = 0) also developed the network to an equilibrium state similar to the linear rule (Fig. 3Dii). However, the Gaussian rule with two non-zero setpoints (*η* = 0.7 and *ɛ* = 7.9) failed to develop a proper network (Fig. 3Diii).

**Fig. 3:**
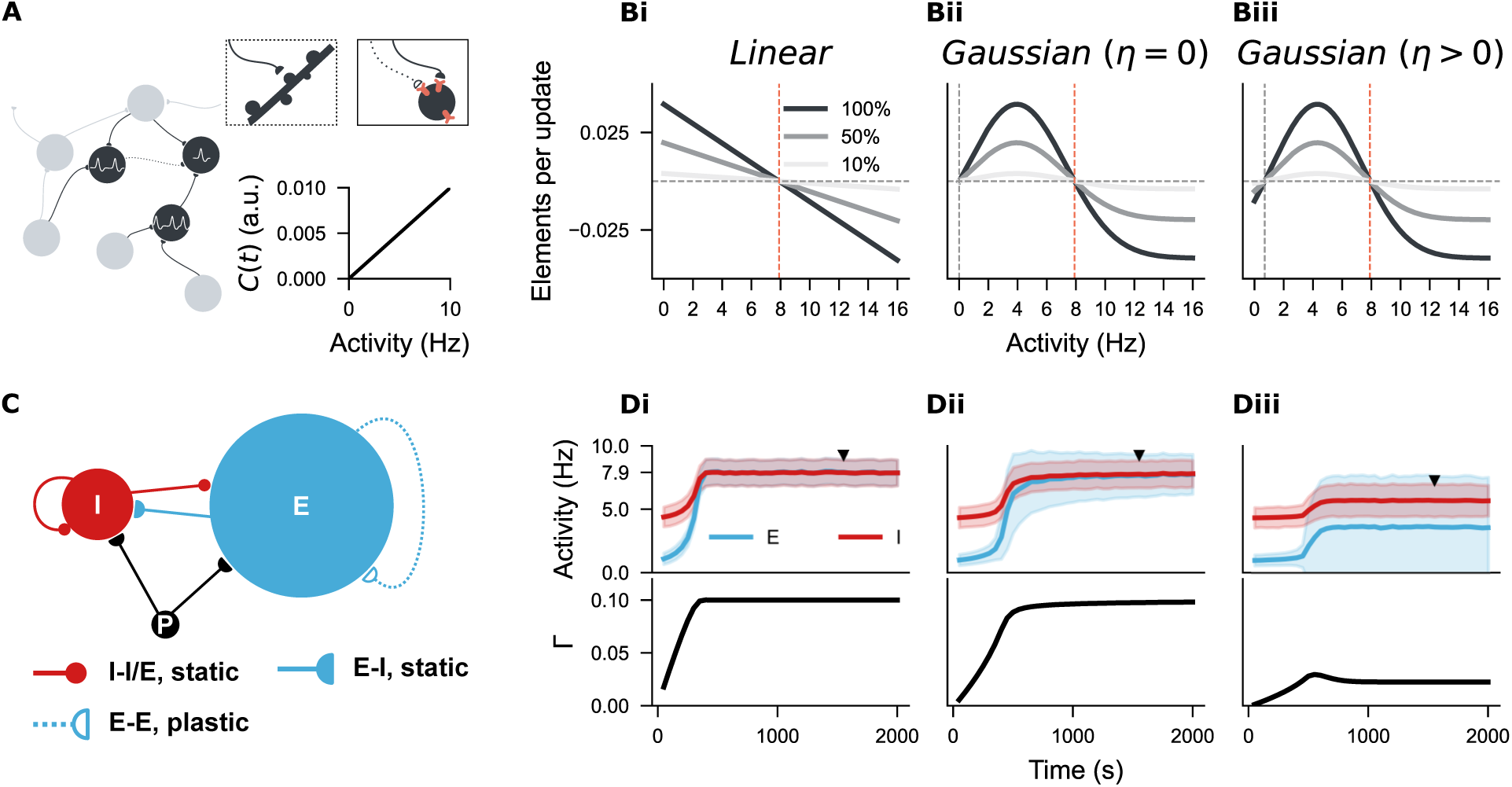
Growing a neural network with three distinct structural plasticity rules. (A) A network of point neurons was used to study structural plasticity, with simplified dendritic morphology. Dendritic spines are depicted as pink sticks on the soma, and axonal boutons are represented by empty or solid half-circles. An empty half-circle with a dashed line indicates an axon during retraction. In this model, intracellular calcium concentration is linearly correlated with neural firing rate, so neural activity is used to reflect “firing rate homeostasis” throughout the manuscript, as it has a meaningful physical unit compared to the arbitrary unit (a.u.) for calcium concentration. (Bi-Biii) Three structural plasticity growth rules regulate changes in synaptic element numbers. (Bi) Linear rule with a single setpoint (*ɛ* = 7.9, orange line). (Bii) Gaussian rule with two setpoints, one at zero (*η* = 0, grey line; *ɛ* = 7.9, orange line). (Biii) Gaussian rule with two non-zero setpoints (*η* = 0.7, grey line; *ɛ* = 7.9, orange line). Three shades represent 100%, 50%, or 10% of the original growth rate (*ν*), with positive and negative values indicating the rate of synaptic element growth or loss. (C) Neural network architecture based on the Brunel network model, consisting of 10 000 excitatory (blue, E) and 2 500 inhibitory neurons (red, I) stimulated by external Poissonian inputs (P). All I-I, I-E, and E-I synapses are hard-wired with 10% probability, while E-E synapses are subject to structural plasticity rules. (Di-Diii) Temporal dynamics of neural activity and network connectivity (Γ) during network growth, guided by the three distinct rules. Unless otherwise stated, the curve and shaded areas in activity plots represent the mean and standard deviation of the neural activity for the I and E populations. The network reached an equilibrium state (Γ = 10%) in Di and Dii but not in Diii. The firing rates distribution and network connectivity matrices of the indicated time points (solid triangles) are provided in Supplementary Fig. 3 for Di-Dii, and in Fig. 4 for Diii.

Closer examination of the neural firing rates and network connectivity at the late growth stage of this network (Fig. 3Diii) revealed that half of the excitatory neurons were silent (Fig. 4A), and the network exhibited inhomogeneous connectivity, with nearly half of the neurons isolated (Fig. 4Bi-Bii). Correlating the firing rates with synapse numbers of individual excitatory neurons confirmed that the silent neurons were isolated, while neurons with firing rates close to the target possessed approximately 1 000 synapses, similar to the linear rule (Fig. 4C). These results align with the properties of the Gaussian growth rules: one setpoint is stable (*ɛ* = 7.9), while the second setpoint can be stable if the synapse numbers remain unchanged after silencing (*η* = 0) or unstable if disconnection occurs after silencing (*η >* 0). We refer to the Gaussian rule with two stable setpoints as *stable Gaussian rule*, while the variant with one unstable setpoint is termed the *biphasic Gaussian rule* in this manuscript.

**Fig. 4:**
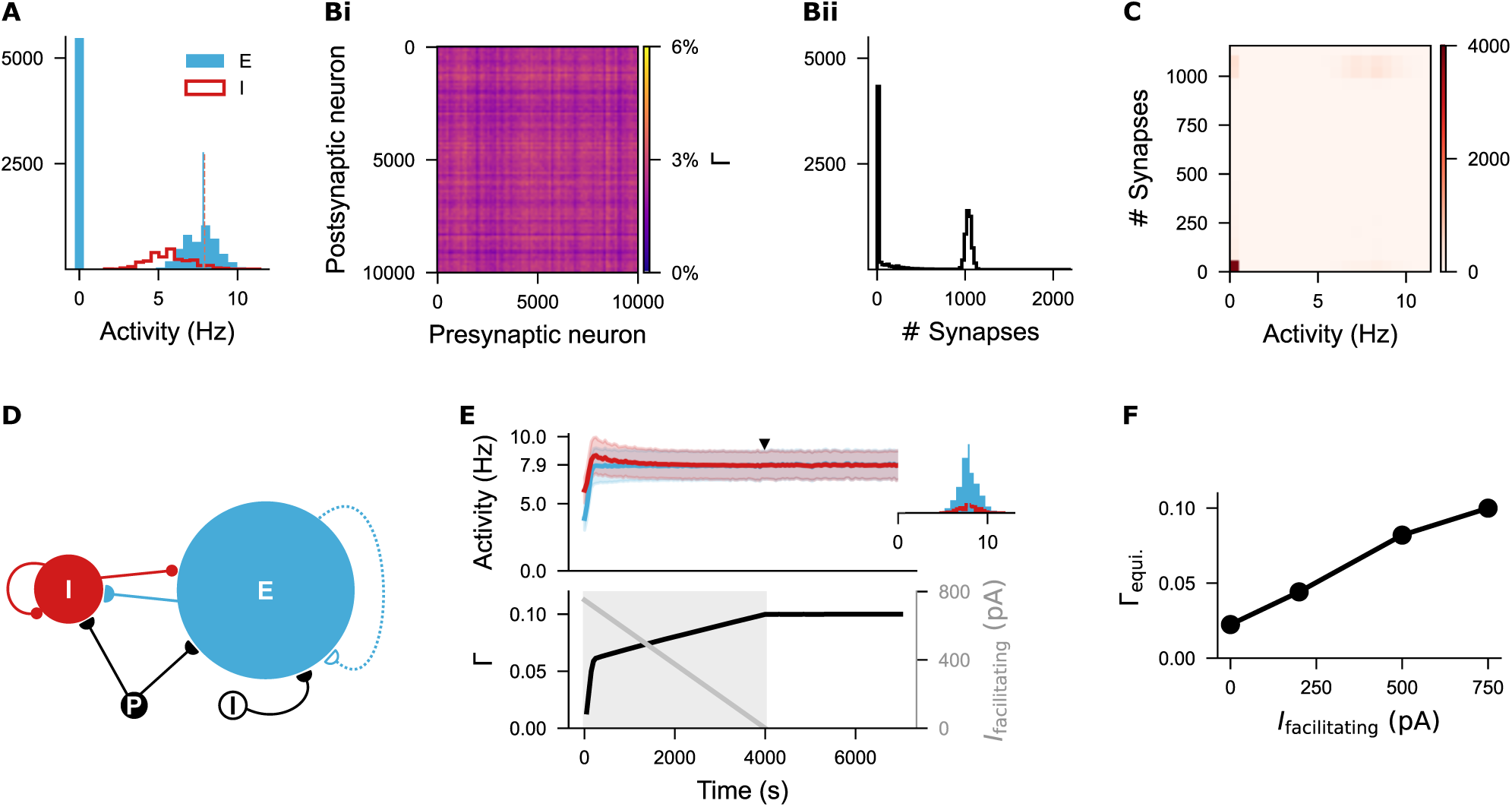
Silent neurons remain isolated in the network regulated by the biphasic Gaussian rule. (A) Histogram of firing rates for excitatory and inhibitory neurons sampled at the time point indicated in Fig. 3Diii. Nearly half of the excitatory neurons were silent. The blue vertical line represents the mean firing rate of non-silent neurons, while the orange dashed line marks the target firing rate (*ɛ* = 7.9 Hz). (Bi-Bii) Network connectivity matrix and distribution of synapse numbers for individual excitatory neurons. (C) Correlation heatmap showing the relationship between neural activity and synapse number of individual excitatory neurons. Silent neurons did not form synapses, whereas active neurons firing around the target rate formed approximately 1 000 synapses with other active excitatory neurons. (D) Network architecture when a damping facilitating current (*I*_facilitating_) was injected to enhance network development. (E) Temporal dynamics of neural activity and network connectivity when a facilitating current was injected. The inset shows the firing rate distributions of both excitatory and inhibitory neurons at a selected time point (solid triangle). The facilitating current started at 750 pA and decayed linearly to zero during a 4 000 s period. (F) Different initial values of facilitating currents resulted in varying network connectivities. An initial value of 750 pA was used throughout the manuscript.

Inspired by previous observation that increased neural excitability facilitates circuit development,^54^ we enhanced the excitability of excitatory neurons during network growth by applying a damping facilitating current (*I*_facilitating_) to raise the average membrane potential closer to the threshold potential (Fig. 4D). With this intervention, the neural network governed by the biphasic Gaussian rule successfully grew to the equilibrium state and maintained the dynamics after the facilitating current decayed to zero (Fig. 4E). Given that varying the initial intensity of the facilitating current resulted in different equilibrium connectivity (Fig. 4F), we used the intensity (750 pA) that produced the same network connectivity (10%) and firing dynamics as the linear rule for further exploration.

### The biphasic Gaussian rule reconciles homeostatic and non-homeostatic structural changes triggered by activity perturbation

As a proof-of-concept, we systematically modulated the input strength to a subgroup of excitatory neurons (10% of the entire excitatory population) after the growth period. Input strength was varied from 0% to 200% fold of the original intensity (FOI, Fig. 5A), mimicking both input deprivation and stimulation. Fig. 5Bi-Biii,C illustrates three example protocols demonstrating neural activity and network connectivity under the biphasic Gaussian rule. Intuitively, stimulation increased the activity of the targeted subpopulation (S) (left panel in Bii) and triggered a reduction in connectivity both among the stimulated neurons (S-S) and between the stimulated and non-stimulated excitatory neurons (S-E; left panel in Biii). The connectivity matrices in panel C display the final network connectivity. Ultimately, homeostatic disconnection restored the activity levels of the stimulated neurons to the setpoint value (orange line in Bii). In contrast, the outcome of activity deprivation depended on its severity. Both mild deprivation and silencing reduced the neural activity of the affected subpopulation (middle and right panels in Bii). However, only mild deprivation allowed the network to restore activity to the setpoint value via a homeostatic increase in network connectivity (middle panels in Biii and C). In the case of complete silencing, the affected neurons disconnected from the network (right panels in Biii and C).

**Fig. 5:**
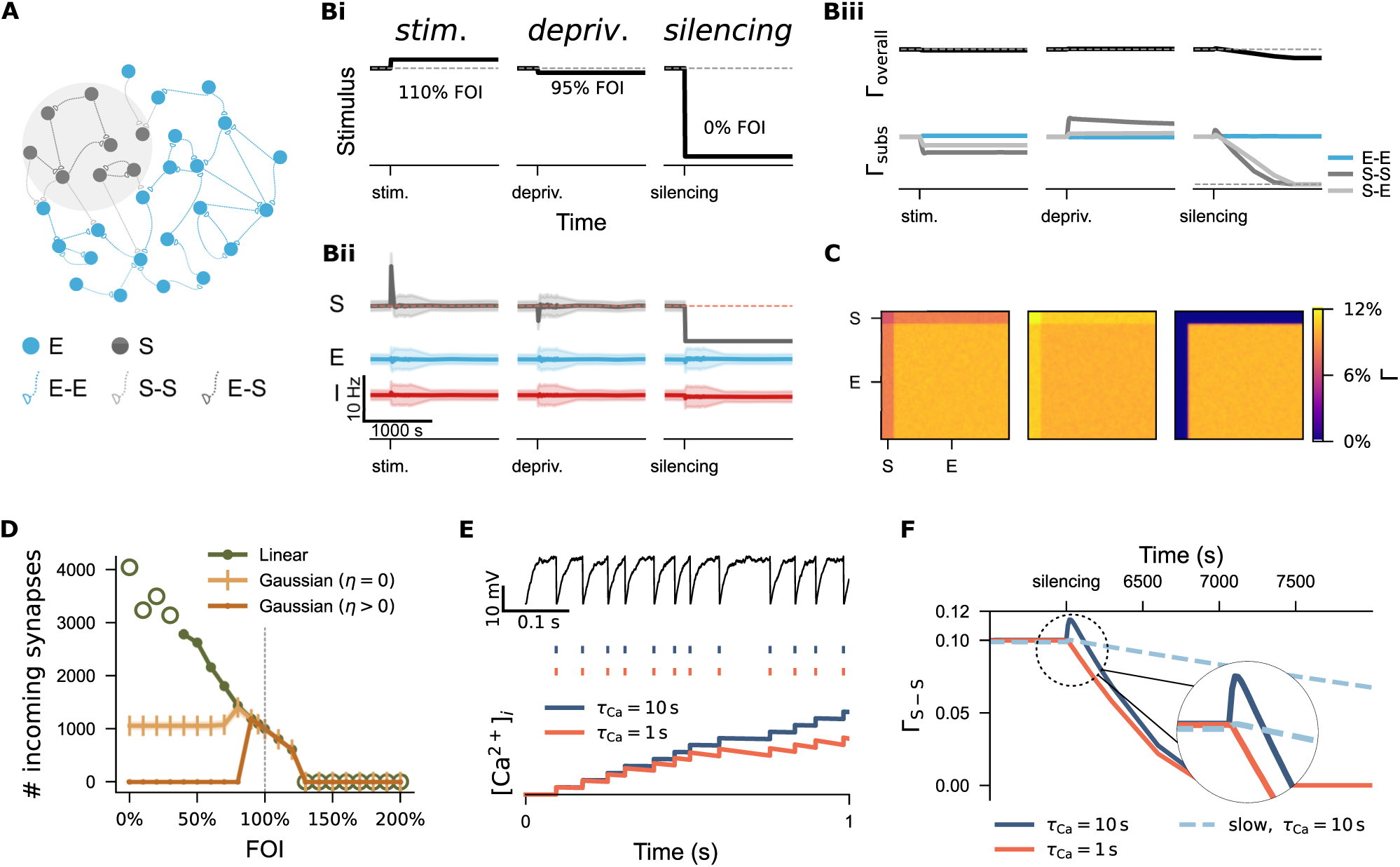
Divergent regulation of network connectivity under stimulation and deprivation via three structural plasticity rules. (A) A subpopulation (S) consisting of 10% excitatory neurons (E) was subject to activity perturbation. All E-E, S-S, and E-S synapses were governed by the biphasic Gaussian rule. (Bi) Activity perturbation protocol. Three different folds of the original intensity (FOI) from the Poisson generator were used as examples to represent *stimulation* (110% FOI), *mild deprivation* (95% FOI), and *silencing* (0% FOI). (Bii) Temporal dynamics of neural activity of S, E, and inhibitory neurons (I) under each protocol. (Biii) Temporal evolution of the overall network connectivity (Γ_overall_) and subgroup connectivity (Γ_subs_) under each protocol. The synaptic connection probability from E to S is identical to that from S to E, as the same rules were applied to spines and boutons. Therefore, only S-E traces are shown unless otherwise stated. (C) Network connectivity matrices at the end of the three protocols. (D) Average incoming synapse numbers of S neurons under different FOIs. The empty green circles represent data from networks subjected to extreme stimulation or inhibition, where both neural activity and network connectivity were unstable. (E) Examples of two neurons receiving the same external inputs but with different calcium decay time constants (*τ*_Ca_). The upper panel shows membrane potential; the middle panel displays spike trains; the lower panel depicts the integrated calcium concentration over time. (F) Connectivity traces of the subnetwork under silencing across three different conditions. The dashed circle area is shown at higher magnification.

A systematic analysis of incoming synapse numbers confirmed the biphasic dependency of the biphasic Gaussian rule (dark yellow curve in Fig. 5D) and the monotonic behavior of the linear rule (green curve). The stable Gaussian rule, which has two stable setpoints (*η* = 0), exhibited an intermediate response: external stimulation and mild deprivation triggered a homeostatic reduction or increase in synapse numbers, but strong deprivation left the network silent and intact (light yellow curve). Of the three rules, the biphasic Gaussian rule best captured the homeostatic properties of the linear rule while also allowing for non-homeostatic, silencing-induced spine loss, consistent with our experimental findings and previous studies.^55–59^

Lesion-or denervation-induced plasticity is of particular interest in activity-dependent structural plas-ticity and has clinical relevance.^60–62^ The biphasic Gaussian rule provides a useful model for studying denervation-induced degeneration and subsequent regeneration. However, we observed a small, biologically unrealistic increase in connectivity immediately following silencing, possibly due to a fast growth rate or residual effects of prior activity on intracellular calcium concentrations. In Fig. 5E, we showed that two neurons receiving the same external inputs displayed identical membrane potential dynamics and spike trains, but difference in the calcium time constant (*τ*_Ca_) caused a divergence in accumulated calcium concentration. Since synaptic element numbers depend on intracellular calcium concentration in our model, which reflects firing rate, a shorter time constant (*τ*_Ca_ = 1 s, orange curve in 5F) carried less activity history, leading to a smoother reduction in synaptic connectivity. A similar effect was achieved by slowing the growth rate (10% of the original rate) for structural plasticity (light blue dashed curve, 5F). Our computer simulations showed that the timescales of calcium dynamics and structural plasticity interact. The decay time constant for calcium concentration is difficult to estimate due to the diversity of calcium signals, which vary in time and spatial compartmentalization. Additionally, structural plasticity is typically a slow process, occurring over minutes, hours, and days. Therefore, we used the slow growth rate and the original calcium time constant for subsequent studies. The faster growth rate employed in Fig. 6D-F was used solely to accelerate spine turnover, thereby reducing simulation time and allowing us to observe long-term changes more quickly.

**Fig. 6:**
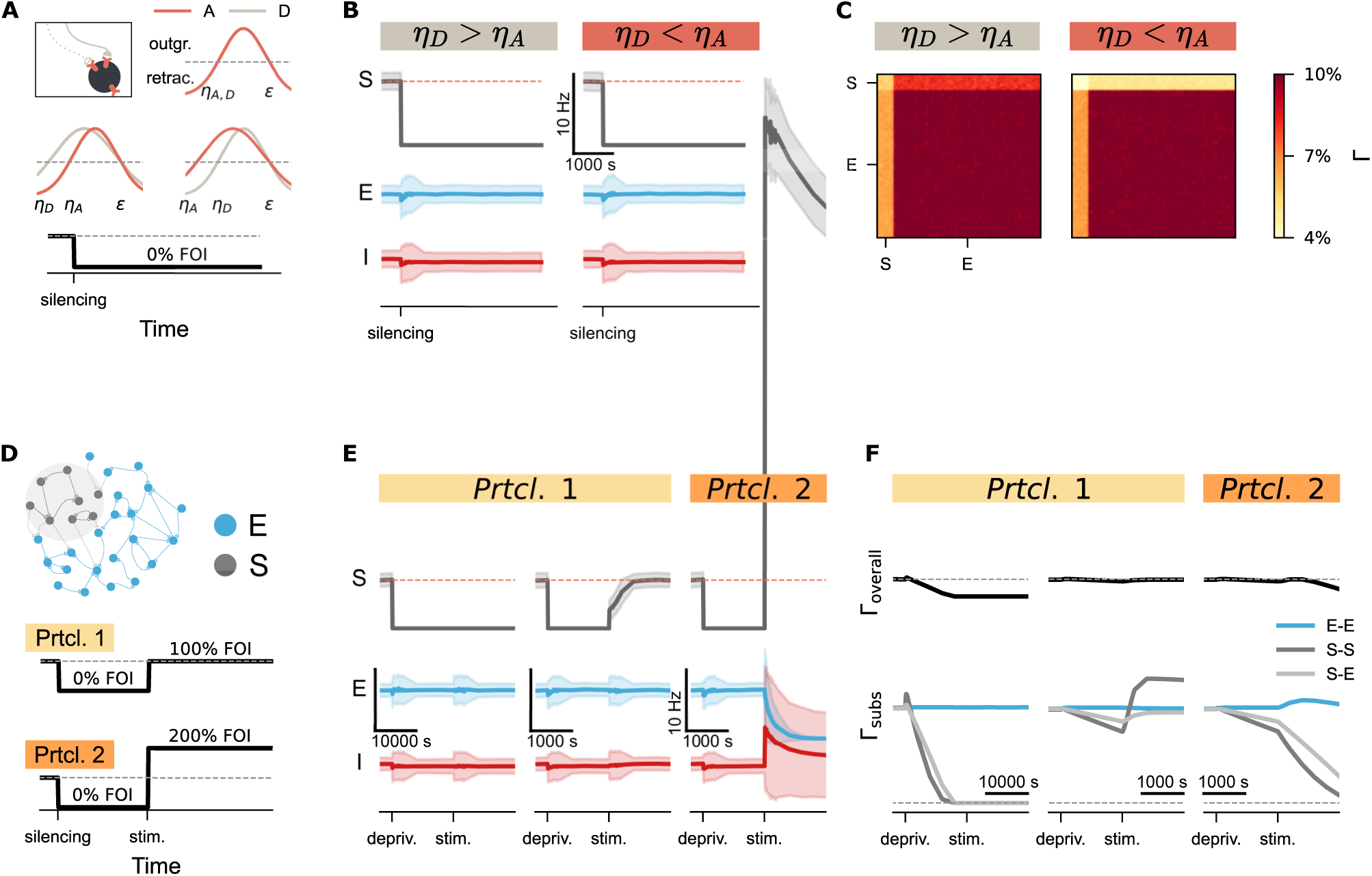
Activity perturbation and recurrent connectivity shape the evolution of network connectivity. (A) In the default network, the same growth rule was applied to both axonal boutons (A, light brown curve) and dendritic spines (D, pink curve), with *η_A_* = *η_D_*, as shown in the upper inset. Different *η* values could be used for axonal and dendritic elements, such as *η_A_ > η_D_* or *η_A_ < η_D_*. A silencing protocol was applied (fold of original intensity, FOI). (B) Neural activity of the stimulated subpopulation (S), excitatory neurons (E), and inhibitory neurons (I) under two conditions. (C) Network connectivity matrices at the conclusion of the silencing protocol under the two conditions. (D) Protocols used to investigate the effects of recurrent connectivity and external stimulation. The same growth rule was applied to both axonal and dendritic elements (*η_A_* = *η_D_*) here. (E, F) Time courses of neural activity and connectivity following silencing and external stimulation. In Protocol 1, the growth rate for the left panel (Prtcl. 1; left panel) was ten times faster than in the left panel (Prtcl. 1; right panel). In Protocol 2 (Prtcl. 2), the intensity of external stimulation was doubled as compared to the one used in Prtcl. 1.

### Recurrent connectivity and activity perturbation shape spine number evolution during synaptic recovery

While silencing-induced disconnection captured the characteristic spine loss observed during input deprivation, complete neural isolation is rarely observed *in vivo*. Instead, spine recovery often occurs following an initial loss of spines.^55, 57, 59^ Previous computer simulations using a Gaussian rule have also reported such “physiological” recovery, assuming distinct growth rules for axonal bouton and dendritic spines.^43, 44^ However, when we applied different *η* values for axonal boutons and dendritic spine growth within a topology-free network (without distance-dependent connectivity), synapse numbers failed to recover after silencing (Fig. 6A,B). Instead, we observed only asymmetric connectivity in the input and output synapses of the deprived subpopulation (Fig. 6C). A closer examination of the network revealed the critical role of recurrent inputs. In networks with distance-dependent connectivity, neurons near the boundary of the deprived region continued receiving active inputs from non-deprived neighbors, making them less deprived than neurons at the center, even though all external inputs were equally removed.

This observation led us to further investigate the role of recurrent inputs in synaptic recovery. In our topology-free network, we assessed the effects of recurrent inputs by adjusting both external inputs and internal connectivity. After applying the silencing protocol, we introduced external stimulation to the deprived subnetwork with different recurrent connectivities. In Fig. 6E,F (Protocol 1), the network in the left panel was simulated at a speed ten times faster than in the middle panel, meaning that the recurrent connectivity was much lower in the left panel when stimulation was applied. To simplify the analysis, we avoided complicating factors related to stimulation intensity – such as stimulation magnitude, duration, interval, and session numbers – that have been studied previously.^19, 49, 63^ Instead, we kept the stimulation constant and used two different intensities as a proof of concept. The results demonstrated that network dynamics are determined by the product of external inputs and internal connectivity. In the left panel of Fig. 6E, the deprived subnetwork remained silent despite stimulation, due to low recur-rent connectivity. In contrast, the same stimulation reactivated the deprived subnetwork in the middle panel, where connectivity was higher. Connectivity traces in Fig. 6F confirmed that residual recurrent connectivity amplified the external stimulation, leading to synapse regeneration and rewiring. However, if the external stimulation was too strong (protocol 2), the amplification effects of recurrent connectivity caused the system to shift from silencing-induced degeneration to over-excitation-induced homeostatic degeneration (Fig. 6E,F, right panels). These simulation results suggested that recurrent connectivity, in combination with external inputs, jointly determines neural activity and influences synapse numbers and network connectivity.

### Synaptic scaling shapes intrinsic connectivity and modulates structural plasticity during activity perturbation

In line with this concept, homeostatic synaptic scaling could theoretically reshape recurrent connectivity and modulate neural activity by dynamically adjusting functional synaptic transmission. Specifically, homeostatic downscaling of excitatory synapses might reduce heightened neural activity, potentially masking changes in structural plasticity, particularly if it operates quickly and effectively. Conversely, homeostatic strengthening of excitatory synapses could increase neural activity. Under conditions of silencing, this strengthening may shift activity-dependent structural plasticity from non-homeostatic, silencing-induced synapse loss to homeostatic rewiring, similar to the outcomes observed with external stimulation or distance-dependent connectivity.

To test this hypothesis, we implemented a monotonic, activity-dependent synaptic scaling rule guided by the same intracellular calcium concentration as the structural plasticity rule. To ensure consistent baseline connectivity and avoid developmental issues, we allowed the network to grow using the biphasic Gaussian structural rule until it reached the equilibrium state. We then activated the calcium-based synaptic scaling at the onset of the activity perturbation protocol (Fig. 7A; Supplementary Fig. 4A). To disentangle the contributions of structural plasticity and synaptic scaling, we analyzed two types of connectivity: (1) *Structural Connectivity* (Γ_struc._), determined solely by the number of synapses; and (2) *Effective Connectivity* (Γ_effec._), calculated by multiplying the number of synapses by the weight of each synapse. Without synaptic scaling, synaptic weights remained uniform, resulting in identical values for structural and effective connectivity (Fig. 7C, left panels; Supplementary Fig. 4Bi). Intuitively, when neural activity was increased through stimulation, both structural plasticity and synaptic scaling operated redundantly in a homeostatic manner, leading to a less pronounced homeostatic disconnection in the presence of synaptic down-scaling.

**Fig. 7:**
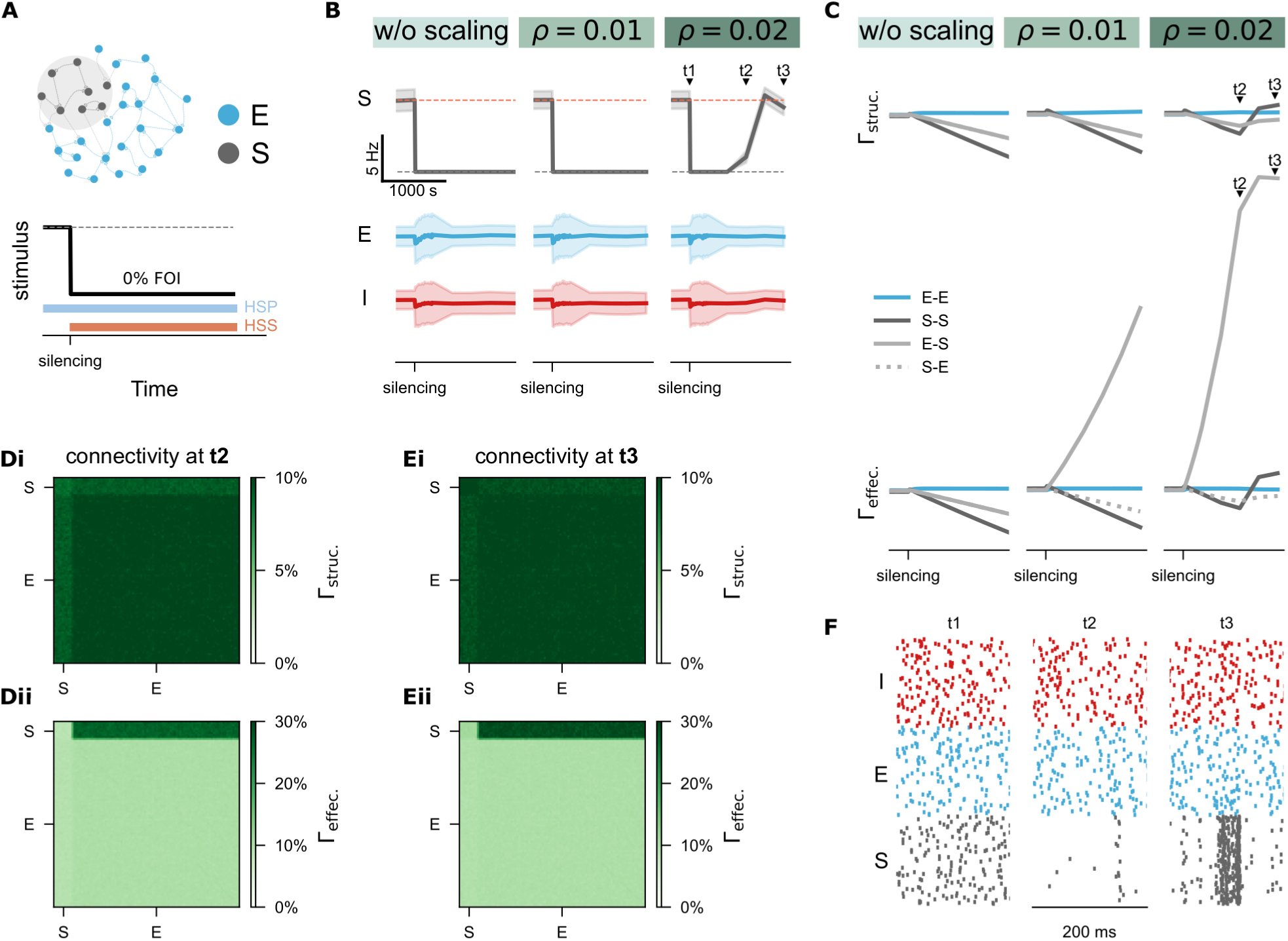
Homeostatic synaptic scaling (HSS) shapes effective connectivity and interacts with homeostatic structural plasticity (HSP). (A) Protocol of silencing – 0% FOI, applied to a subpopu-lation of excitatory neurons (S) – and synaptic scaling activation. Three different scaling strengths were applied: *ρ* = 0 represents no scaling, *ρ* = 0.01 represents weak scaling, and *ρ* = 0.02 represents strong scaling. (B-C) Time courses of network activity and connectivity. Γ_struc._ refers to synapse-number-based structural connectivity, while Γ_effec._ represents effective connectivity, which is the product of synapse numbers and synaptic weights. (Di-Dii) Structural and effective connectivity matrices of the entire net-work at *t*2. (Ei-Eii) Structural and effective connectivity matrices of the entire network at *t*3. (F) Raster plots for 100 selected neurons, including inhibitory (red, I), excitatory (blue, E), and silenced excitatory neurons (grey; S), from the network at the time points *t*1, *t*2, and *t*3.

In contrast, activity deprivation involved a complex interplay between synaptic scaling and structural plasticity. With weak scaling (Fig. 7B,C, *ρ* = 0.01), there was a gradual increase in effective connectivity to the deprived neurons (E-S), but this had minimal impact on firing rates and was insufficient to restore synapse numbers and network connectivity. Increasing the scaling strength (*ρ* = 0.02), as shown in the right panel of Fig. 7C, resulted in a more pronounced increase in effective connectivity in E-S synapses, which successfully reactivated the deprived neurons from their silent status, initiating synapse regeneration and rewiring. We also examined the structural and effective connectivity matrices at time points *t*2 (Fig. 7Di,Dii) and *t*3 (Fig. 7Ei,Eii). Notably, while the number of input synapses to the deprived subgroup decreased at *t*2, the remaining synapses that received inputs from the non-deprived neurons increased in synaptic weight. This reactivated the silent neurons (Fig. 7B, right panels), and by *t*3, the number of input synapses to the deprived subgroup began to recover. This observation confirms that monotonic synaptic scaling operates in opposition to the biphasic structural plasticity rule under conditions of severe activity deprivation, facilitating a transition from non-homeostatic synapse loss to homeostatic rewiring. Intriguingly, with sufficient scaling strength, synapse numbers and average neural activity were restored following silencing, and the temporal dynamics of the network were also altered after synapse regeneration. The activity pattern of the previously deprived population showed high synchronization after recovery (Fig. 7F), highlighting mechanisms that may be relevant to post-traumatic conditions such as injury-triggered epilepsy. Overall, our simulation results suggested that biphasic homeostatic structural plasticity and monotonic homeostatic synaptic scaling serve complementary and redundant roles in maintaining firing rate homeostasis, providing a biologically plausible explanation scenario through their interplay.

### Spine size dynamics suggest an interaction between structural plasticity and synaptic scaling

The altered dynamics in the deprived neural population after recovery may result from dominant inputs from non-deprived excitatory neurons. To test this, we analyzed the firing rates and input synaptic weights of representative neurons from both the deprived (grey) and non-deprived (blue) excitatory populations (Fig. 8A). This analysis revealed a notable increase in synaptic weights from non-deprived to deprived neurons (Fig. 8A, lower panel), which was further corroborated by the temporal changes in synaptic weight distribution. Additionally, we assessed input synapse numbers for the representative neuron (solid line) and its peers (dashed line in Fig. 8C) to gain insights into structural plasticity. Consistent with experimental findings, the input synapse numbers of the deprived neurons decreased following silencing but gradually recovered. Interestingly, recovery patterns differed: the synapse numbers from reactivated neurons exceeded initial values (upper panel), whereas those from non-deprived neurons (middle panel) and the total synapse counts (lower panel) restored to slightly lower levels than before the perturbation. Our simulation results demonstrated that an appropriate combination of synaptic scaling and structural plasticity can replicate the process of spine loss and recovery following activity deprivation. Importantly, the model aligns with the commonly observed outcome of chronic activity silencing: fewer but larger-weight synapses, corresponding to fewer but larger spines.

**Fig. 8:**
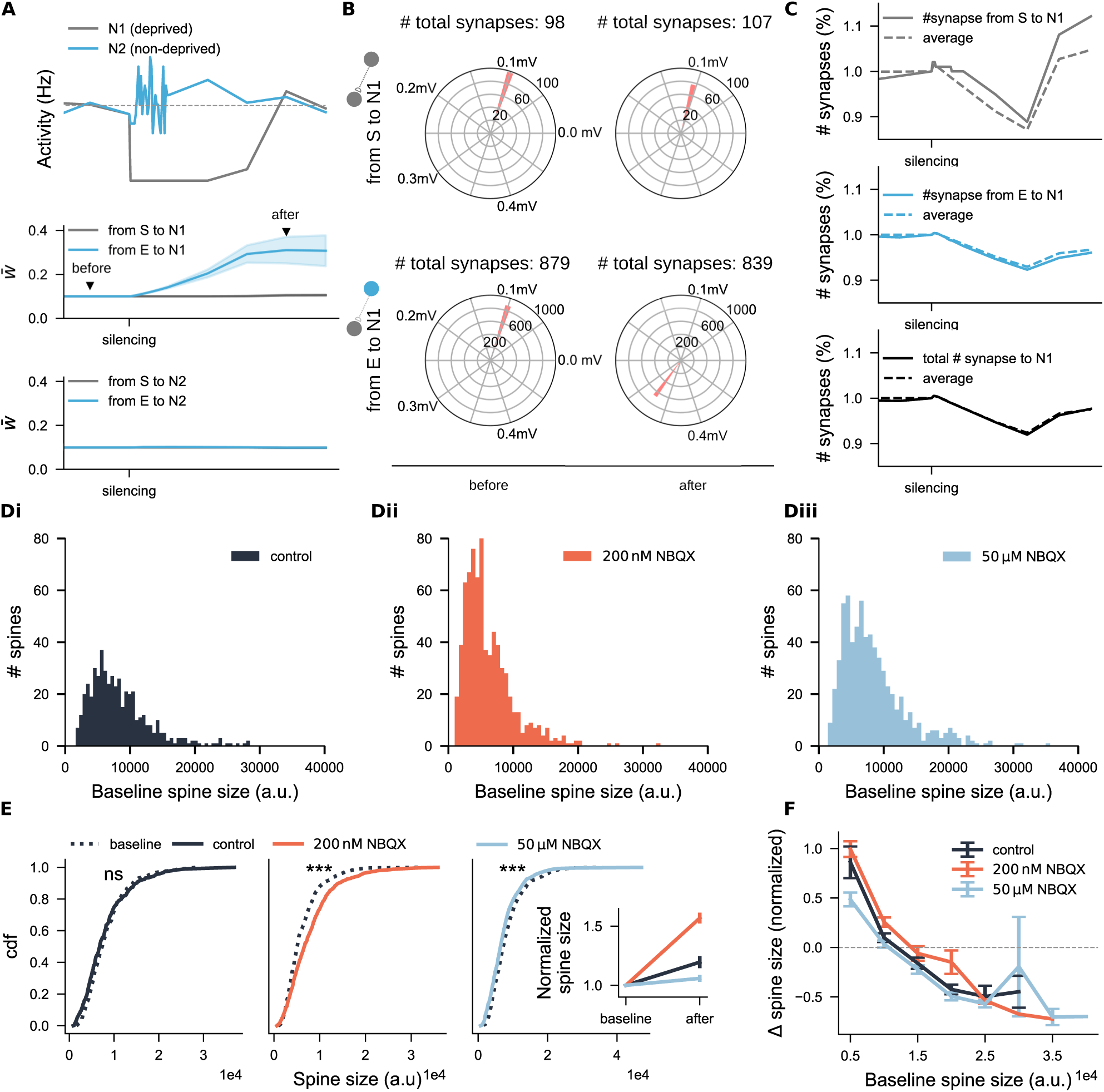
Spine size analysis suggests interactions between functional and structural plasticity. (A) Activity trajectories and average weights of incoming excitatory synapses to two sample excitatory neurons, N1 (deprived) and N2 (non-deprived). The grey dashed line in the upper panel indicates the setpoint activity level. Incoming synapses were grouped into two categories: those from the deprived subpopulation (S, grey) and those from non-deprived excitatory neurons (E, blue). The triangles in the middle panel mark the time points at which synaptic weight distributions were analyzed in panel B. (B) Synaptic weight distributions of neuron N1 at two time points, labeled as *before* and *after*. The numbers indicate the total synapse count for each type at the corresponding time point. (C) Normalized synapse number of the neuron N1 (solid line) and the averaged value across all deprived neurons (dashed line). (Di-Diii) Baseline spine size distributions for three groups. (*N* = 489 for the control group, *N* = 736 for the 200 nM-NBQX group, *N* = 675 for 50 *µ*M-NBQX group) (E) Cumulative distribution function (cdf) of spine sizes before and after the three-day treatment. The inset shows the corresponding averages of normalized spine sizes. (F) Normalized changes in spine sizes, grouped by their initial spine sizes. The values on the *x*-axis represent the upper limits of each binning group. Data points above the dashed line indicate spine enlargement, while those below indicate spine shrinkage.

To verify if this principle holds in our NBQX experiment and to explore any dose dependency, we analyzed spine sizes before and after the chronic NBQX treatment in entorhinal-hippocampal slice cultures. Initially, spine sizes in all three groups showed a long-tail distribution, with a predominance of small spines (Fig. 8Di-Diii). After the three days of treatment, changes in spine sizes (Δ spine size) followed a normal distribution (Supplementary Fig. 3Bi-Biii). At the population level (Fig. 8CD), significant increases in spine sizes were observed following 200 nM (*p <* 0.001, Wilcoxon test; 99.9%CI = [9.09, 25.8], LLM), while 50 *µ*M NBQX treatment lead to significant reductions (*p <* 0.001, Wilcoxon test; 99.9%CI = [9.09, 25.8], LLM), suggesting a non-monotonic, activity-dependent response. When normalizing the size changes by initial spine size, we observed a general increase across all three groups (Fig. 2D inset), likely driven by the enlargement of many small spines. Sorting the spine size changes by initial sizes (Fig. 2E) showed that in control segments (black curve), small spines generally grew while large spines shrank over the three-day period. Under partial inhibition with 200 nM NBQX (orange curve), the dynamics favored an overall increase in spine sizes, regardless of initial sizes. However, with complete inhibition using 50 *µ*M NBQX (light blue curve), only a small number of large spines increased in size, counteracting the natural growth of small spines and causing medium-sized spines to shrink.

These spine data suggested a general homeostatic adjustment of spine sizes in response to partial and complete inhibition. Notably, under complete inhibition, the spine size changes were dependent on initial spine sizes, mirroring experimental findings and our simulation results, where complete activity blockade resulted in fewer but enlarged spines. However, this pattern does not fully align with synaptic weight changes described by the synaptic scaling rule. This discrepancy will be further explored in the discussion, where we integrate electrophysiological and imaging data to provide a comprehensive interpretation.

### Hybrid combinations of synaptic scaling and structural plasticity resolve spine density inconsistencies

Having obtained both spine density and size data for individual segments, we explored whether a universal rule might link spine density and size changes across dendritic segments. We quantified the average spine size changes (Δ̅ spine size) for each segment and correlated these with baseline spine densities (Fig. 9A) and changes in spine density (Δ spine density, Fig. 9B). A significant positive correlation between spine size changes and baseline spine densities was observed in the 200 nM NBQX treatment group (Fig. 9A, *r*(24) = 0.65, *p* = 0.00064, Pearson’s correlation; Supplementary Fig. 2A), but this correlation was absent in the control and 50 *µ*M NBQX groups. When correlating average spine size changes with alterations in spine density across all segments, weak inhibition (200 nM NBQX) typically resulted in increases in both spine density and size in most segments (see orange dots clustered in the upper right quadrant in Fig. 9B). This suggested that spine-dense segments may play a key role, with a synergistic interaction between structural and functional plasticity under weak inhibition. In contrast, under strong inhibition with 50 *µ*M NBQX, the size-density data points were more widely scattered across the four quadrants (light blue dots in Fig. 9B), reflecting the diverse spine size modifications observed. Although our imaging data did not reveal a universal rule applicable to all segments, the results suggested a highly variable response to activity deprivation across individual dendritic segments. This led us to hypothesize that different combinations of synaptic scaling and homeostatic structural plasticity may operate at the dendritic segment level or even vary across different cell types (Fig. 9C). This diversity may explain the inconsistent findings observed in neurons from different regions or the contrasting effects seen between apical and basal dendrites within the same neuron.^64^

**Fig. 9:**
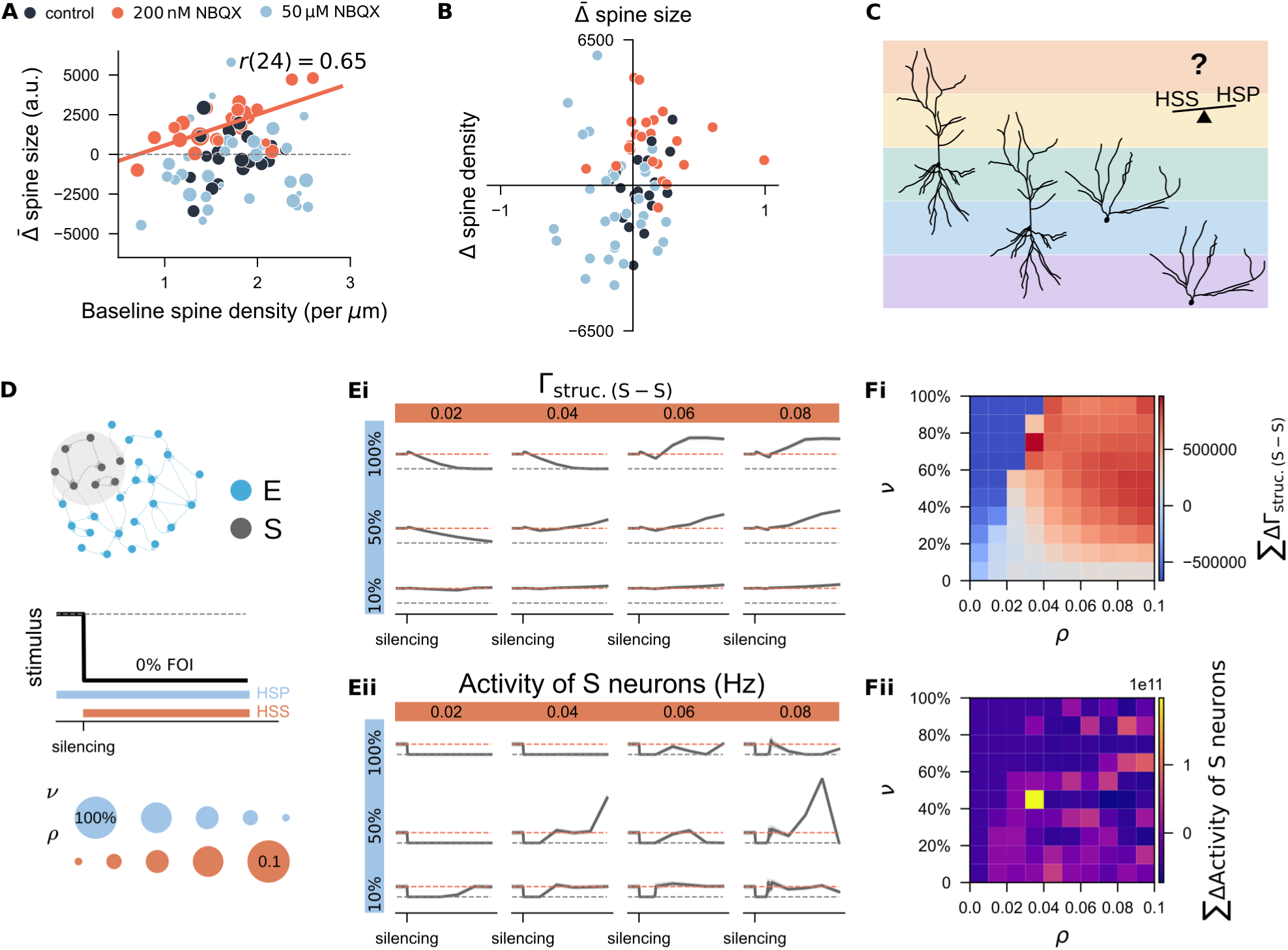
Systematic study of the interaction between synaptic scaling and structural plasticity in response to activity silencing. (A) Correlation between each dendritic segment’s average change in spine size and its initial spine density. Marker size represents the net change in spine density over a three-day period. (B) Correlation between each dendritic segment’s average changes in spine size and its change in spine density. (C) Hypothesis: Different combinations of homeostatic synaptic scaling (HSS) and homeostatic structural plasticity (HSP) may form a spectrum, allowing different neuron types or dendritic segments within the same neuron to be governed by unique subsets of these rules. This spectrum could explain the high diversity observed in empirical studies of structural plasticity. (Sample neurons were reconstructed based on a CA1 pyramidal neuron (left two) and a dentate gyrus granule cell (right two), previously recorded in our lab.) (D) Simulation protocol. We systematically varied the growth rate of the HSP rule (*ν*) and the scaling strength of the HSS rule (*ρ*). (Ei-Eii) Example traces of structural connection probability and neural activity of the deprived subpopulation (S) under different parameter combinations. (Fi-Fii) Discrepancies in connectivity and firing rates when different growth rates and scaling strengths were combined. Discrepancies were calculated by estimating the area between the actual time course and the equilibrium connection probability (10%) or the target rate (*ɛ*) from the time of silencing until the end of the simulation. Results are averaged from 11 random trials and displayed in panels E and F.

To explore this hypothesis, we applied the same silencing protocol in simulation, varying the growth rates of structural plasticity (*ν*) alongside different synaptic scaling strengths (*ρ*) (Fig. 9D). Example trajectories of neural activity and structural connectivity in the deprived subpopulation under different conditions are shown in Fig. 9Ei,Eii. To evaluate the effect of these parameter combinations on network rewiring, we summed the differences between actual structural connectivity in the subnetwork and the equilibrium state value (10%) over the entire silencing period (Fig. 9Fi). Cold colors represent a failure in connectivity restoration, while warm colors indicate successful rewiring or over-reconnection. Obser-vationally, slower growth rates led to gradual spine loss, resulting in more residual synapses. In this case, even weak synaptic scaling was sufficient for synapse recovery. In contrast, faster growth rates, associated with fewer residual synapses, required stronger synaptic scaling for comparable restoration.

We also tracked discrepancies between actual neural rates and the target rate (7.9 Hz) for the deprived population over the same period (Fig. 9Fii). Parameter combinations that failed to restore network connectivity (Fig. 7Fi) showed negative deviations from the target rate (dark blue in Fig. 7F), indicating that the reactivation of silent neurons was unsuccessful. However, successful synapse regeneration (warm colors in Fig. 7Fi) did not always align with smooth firing rate restoration. Optimal firing rate restoration was observed with parameters leading to moderate synapse regeneration (light pinkish colours in panel Fii). Combinations of rapid structural growth and strong synaptic scaling could lead to excessively high neural activity (bright colors), mimicking epileptiform activity, or result in decreased transient firing rates (dark blue colors). In conclusion, our systematic study confirmed that calcium-based synaptic scaling and homeostatic structural plasticity interact across varying time scales and strengths. Their interplay helps reconcile the disparate experimental findings reported in previous studies.^50^

## Discussion

The mammalian brain, a complex system with billions of neurons and non-neuron cells organized into layers, regions, and circuits,^65^ is highly robust such that flooding information does not cause catastrophic forgetting,^66^ nor will input deprivation destroy the network dynamics.^15–17^ Hebbian plasticity and homeostatic synaptic scaling (HSS), both well-studied forms of functional plasticity, are central to maintaining this neural network’s functional robustness and dynamics. However, structural plasticity has been relatively undervalued due to inconclusive experimental evidence^50^ and limited computational models.^42^ Under activity deprivation, both homeostatic and non-homeostatic changes in spine numbers have been observed,^50^ posing a challenge to the promising concept of the homeostatic structural plasticity (HSP) model. This manuscript details the biphasic HSP model and its interaction with the monotonic HSS under activity deprivation. Our time-lapse imaging experiments confirmed the non-monotonic activity dependency of spine numbers, supporting the biphasic HSP rule. Simulations demonstrated that this rule reconciles homeostatic and non-homeostatic changes, complementing the HSS rule in adjusting synaptic weights and influencing synapse numbers. The redundancy and heterogeneity between HSP and HSS rules are essential for maintaining firing rate homeostasis and ensuring the stability and functionality of brain networks.

Our research significantly expands the understanding of structural plasticity by providing evidence of its differential, activity-dependent regulation. Although structural plasticity is often associated with functional cortical reorganization following injury and amputation or in cases of disease-related cognitive decline, its modeling has been less extensive compared to functional plasticity due to the challenges of capturing high-dimensional morphological changes. Among the existing mathematical implementations, some focused on dendritic tree development,^67–69^ while others emphasized synapse formation and rewiring as the key differentiator from functional plasticity models. However, these early models mostly adopted a synapse-specific perspective akin to Hebbian plasticity, emphasizing used-based growth and removal of individual synapses,^70–74^ sharing limitations in stabilizing network dynamics. In contrast, Butz and van Ooyen introduced a cell-autonomous HSP rule, rooted in van Ooyen and van Pelt’s work^75^ and aligned with concepts proposed by Dammasch.^76^ This rule employs a setpoint firing rate or calcium concentration to guide spine and bouton formation or elimination.^43, 77^ This spine-number-based rule is a structural analog to synaptic scaling in maintaining firing rate homeostasis. In the meantime, it retains the associative properties of correlation-based models^19, 46, 47^ and promotes self-organized criticality in developing neural networks.^78^ Therefore, in linear or Gaussian format, this rule has long been suggested as an alternative solution to integrate Hebbian and homeostatic plasticity, replicating various phenomena observed in development and disease-like conditions. Our work supports the biological plausibility of a biphasic HSP rule, substantiating it with experiments and computational evidence.

We used gradual activity blockade and time-lapse imaging to propose that spine numbers follow a biphasic relationship with neural activity, addressing inconsistencies in spine density findings across various studies.^50^ Experimental paradigms, such as lesion methods and pharmacological treatments, often introduce confounding factors like neuroinflammatory cytokine production,^57, 58, 79, 80^ which influences plasticity.^81–83^ Non-traumatic deprivation methods, such as dark rearing^84^ or whisker trimming,^85^ result in non-uniform input deprivation across the dendritic tree, complicating interpretations of spine density changes across different dendritic compartments. Precise pharmacological treatments, such as those targeting pre-synaptic mechanisms^86^ or using glutamate uncaging near selected spines,^87^ also have limitations due to the homogeneity of their effects. Even studies using drugs that uniformly target postsynaptic neurons report inconsistent results in spine density,^88–93^ which may have suggested the potential role of the strength of deprivation. Factors like drug type (non-competitive or competitive), experimental setting (*in vivo* or *in vitro*), and circuit integrity significantly influence the degree of inhibition experienced by target neurons. Our recent findings in CA1 pyramidal neurons revealed distinct cellular, structural, and molecular responses to synaptic input deprivation depending on the pathway involved, highlighting the pathway-dependent nature of these changes.^94^ Based on these observations, we hypothesized a bi-phasic activity-dependent rule to explain the wide range of spine density changes: partial activity suppression promotes spine growth, while complete inhibition results in spine loss. Indeed, our time-lapse imaging confirmed this hypothesis, showing increased spine density with partial inhibition (200 nM NBQX) and decreased density with complete inhibition (50 *µ*M NBQX). However, the results also pointed to complex behaviors in spine size upon inhibition, suggesting an interaction between functional and structural plasticity, which we will explore further in the discussion.

The biphasic structural plasticity model, in the shape of a Gaussian growth curve, successfully reconciled the diverse activity-dependent changes of spine density observed in experimental settings in our simulations. With an unstable setpoint (*η >* 0), this model predicts variable outcomes in synapse numbers (corresponding to spine numbers)–ranging from increases to no changes or reductions–following activity perturbation. Notably, synapse loss induced by silencing falls outside the predictive scope of linear (monotonic) models but is captured by the biphasic rule. As a derived feature, this biphasic rule enables phase transition by modulating neural activity through external input or intrinsic connectivity adjustments. For example, timely and appropriately scaled external stimulation can reverse silencing-induced synapse loss, leading to homeostatic rewiring. This insight could inform non-invasive brain stimulation techniques aimed at preventing the negative effects of stroke-induced neural lesions.^95^ Additionally, the homeostatic response of synaptic scaling promotes recovery of synaptic connections, consistent with empirical observations of synapse numbers and weights. This *natural* rewiring process has also been observed in studies by Butz and van Ooyen using the same Gaussian HSP rule,^41, 43, 44^ where they applied distance-dependent connectivity and emphasized the importance of using distinct growth rules for axonal boutons and dendritic spines. Our successful replication of this rewiring in a topology-free homogeneous network supports their findings, highlighting the importance of intrinsic connectivity in regulating synapse loss and network rewiring while freeing the network from architectural constraints. However, our co-simulation of functional and structural plasticity rules revealed a critical yet synergistic interplay between these different forms of plasticity. This interaction, often overlooked, is essential for interpreting experimental data or modeling biological systems. Moreover, inhibitory plasticity may contribute to the physiological recovery of homeostatic structural plasticity, as suggested by numeric studies,^96^ though this was beyond the scope of our study and warrants future exploration.

In this context, we examined how the tandem behavior of homeostatic structural plasticity and functional synaptic scaling shapes our interpretations. Our experiments revealed an increase in both spine size and density following three days of 200 nM NBQX treatment. This enlargement of spine size likely increased the detectability of small spines, suggesting that the observed increase in spine density results from a combination of spinogenesis, spine stability, and the potentiation of smaller spines. Similar confounding factors affect the interpretation of synaptic scaling results. For instance, Turrigiano et al.^26^ observed changes in miniature EPSC amplitudes but not the frequencies after chronic inhibition and excitation, inferring synaptic weight scaling. However, other studies have reported changes in both amplitudes and frequency. The transformation of silent synapses into active ones^97^ can increase the frequencies but lower the average EPSC amplitudes due to smaller synaptic weights. These represent synapse-number-based structural plasticity, which may not always be detectable with conventional imaging techniques, depending on factors like laser intensity or exposure time. Additionally, under complete inhibition, we observed distinct size changes in small and large spines: smaller spines tend to shrink and disappear, while larger spines enlarge, leading to increased sEPSC amplitudes. This could result in different or even contradictory results between electrophysiological recording and imaging experiments. Therefore, spine numbers and synaptic weights may represent two facets of a broader concept–“effective transmission”–and focusing on only one aspect risks missing the complete picture.

Our systematic study successfully replicated the empirical observation that chronic activity inhibition often leads to spine loss, with remaining spines exhibiting enlarged heads.^90, 91^ This phenomenon can be attributed to the shared calcium signaling mechanism underlying both HSP and HSS rules. While the HSP rule consistently incorporated activity-based calcium concentration as a setpoint, most HSS models focus on firing rate setpoint,^98–101^ synaptic weight normalization,^102^ or presynaptic regulation.^103, 104^ Notably, one model proposed an integral feedback control algorithm,^51^ using a variable with an exponential kernel to track neural spiking activity and adjust synaptic weights. At the time, the role of calcium concentration as the key variable tracking neural activity was not yet known.^105^ Leveraging this model, we reinterpreted the internal status using a calcium-based variant to explore the interaction between the HSS and the HSP rules. As a result, our study also served as a thought experiment (Gedankenexperiment) to understand the interplay between functional and structural plasticity. If both are redundantly monotonic, the rule with the faster timescale would dominate. However, when a biphasic HSP rule interacts with a monotonic HSS rule, bifurcation occurs under strong inactivity, as confirmed by our simulation, resulting in neurons with reduced spine density and enlarged spine heads. From an energy-efficient perspective, synapse formation and spine growth are constrained by protein availability and ATP consumption. Thus, it is more efficient for neurons to potentiate existing synapses rather than forming new ones. The combined effects of synaptic scaling and structural plasticity in maintaining network activity are supported by recent theoretical work at the dendritic trafficking level.^106^ Our study highlights the interdependency between these two forms of plasticity, which may explain the divergent results obtained across different brain regions, neuron types, dendritic segments, and experimental settings.

While our study addressed both synaptic scaling and homeostatic structural plasticity, it is important to note that the use of NBQX is not typical in experiments focused on synaptic scaling. NBQX, a competitive antagonist of AMPA receptors, allows for precise modulation of network activity through dose control,^107^ offering greater flexibility compared to tetrodotoxin (TTX), which blocks action potentials by inhibiting sodium channels. This makes NBQX an effective tool for modulating network activity, as demonstrated by our electrophysiological recordings and supported by other *in vivo* and *in vitro* studies.^107, 108^ However, the specific affinity of NBQX for GluA2-containing AMPA receptors, which play a key role in both TTX-induced synaptic scaling^109^ and glutamate-induced spine protrusion in the presence of TTX,^110^ warrants careful consideration. This selective binding raises questions about the broader implications and mechanisms of NBQX action, particularly in experiments where TTX is absent. While no definitive evidence suggests that NBQX alone impairs the synthesis or insertion of AMPA receptors, further research is needed to fully understand its effects on synaptic scaling and structural plasticity.

In summary, the use of calcium signal in both HSP and HSS rules resembles an integral feedback controller (Supplementary Fig. 5), a widely used engineering approach to achieve robust performance despite external perturbations. In both rules, calcium concentration serves as the control signal to drive connectivity kernel modification via negative feedback, which integrates neural firing rates over time and returns a filtered signal responding to its activity. However, the HSP and HSS rules differ in their detailed mechanism. The HSP rule more closely aligns with a Proportional-Integral-Differential (PID) controller, while the HSS rule operates more like a Proportional-Integral (PI) controller. Besides sharing the integral (I) component, the HSP rule’s proportional (P) component is linked to the growth rate of synaptic elements,^45^ which modifies the number of synaptic elements created or deleted per time unit based on a magnitude relative to the difference between the expected and actual calcium concentration. The scaling factor (*ρ*) in the HSS rule regulates the proportional effects of synaptic scaling. Uniquely, the HSP rule includes a derivative (D) component, determined by the steepness of the growth curve, which is determined by the steepness of the growth curve and dictates how quickly or slowly the system reaches the desired neural activity. This derivative component is critical for ensuring a smooth and robust response to stimuli, avoiding biologically unrealistic overshooting and abrupt oscillations. This is particularly relevant in the Gaussian case, where the distance to the target firing rate accelerates differently depending on the position on the calcium concentration axis. Thus, although the exact shape of the growth rule for homeostatic structural plasticity is still uncertain, we propose that a biphasic curve with a changing slope provides a redundant and heterogeneous structural backup for synaptic scaling, helping maintain robust firing rate homeostasis. This perspective also helps anticipate interactions with other plasticity rules where sub-or supra-threshold calcium dynamics are involved.^111, 112^

## Materials and Methods

### Ethics statement

We used mouse pups postnatal at 3 to 5 days (P3-P5) from C57BL/6J (wild-type) and Thy1-eGFP mouse lines to prepare entorhinal-hippocampal tissue cultures in the current study. All animals were kept under a 12 h *−*12 h light-dark cycle with food and water provided *ad-libitum*. One male and one or two female(s) were kept within the same cage for breeding. All animal experiments were approved by the appropriate animal welfare committee and the animal welfare officer of Albert-Ludwigs-University Freiburg, Faculty of Medicine under X-21/01B and X-18/02C. All effort was made to reduce the pain or distress of animals.

### Preparation of tissue cultures

Entorhinal-hippocampal tissue cultures were prepared as published before.^113, 114^ All tissue cultures were cultivated for at least 18 days inside the incubator with a humidified atmosphere (5% CO_2_ at 35 *^◦^*C to reach an equilibrium status. The incubation medium consists of 50% (v/v) 1*×* minimum essential media (#21575 *−* 022, Thermo Fisher, USA), 25% (v/v) 1*×* basal medium eagle (#41010 *−* 026, Thermo Fisher, USA), 25% (v/v) heat-inactivated normal horse serum, 25 mM 1 M HEPES buffer solution (#15630 *−* 056, Gibco), 0.15% (w/v) sodium bicarbonate (#25080 *−* 060, Gibco), 0.65% (w/v) glucose (#RNBK3082, Sigma), 0.1 mg*/*ml streptomycin, 100 U*/*ml penicillin, and 2 mM glutamax (#35050 *−* 061, Gibco). The incubation medium was renewed 3 times per week. If not stated otherwise, the incubation medium applied to the tissue cultures was always pre-warmed to 35 *^◦^*C and adjusted around pH = 7.38.

### Experimental design

The main objective of the experiments in the current study was to probe whether there is a linear or non-linear dose-dependent regulation of dendritic spine density such that we can implement a spine-number-based structural plasticity rule close to biological reality for further systematic studies. Competitive AMPA receptor antagonist NBQX (2,3-dioxo-6-nitro-7-sulfamoyl-benzo[f]quinoxaline) at different concentrations was bath applied to wild-type cultures while recording synaptic transmission at CA1 pyramidal neurons. Two magnitudes of inhibition (“partial” and “complete”) were thus determined by the reduction of amplitudes and frequencies of spontaneous excitatory postsynaptic currents (sEPSCs). Then the same concentration ladders were used to treat Thy1-eGFP cultures for three days, where we tracked individual dendritic segments of CA1 pyramidal neurons before and after the three-day treatment to investigate whether two different magnitudes of AMPA-receptor inhibition resulted in divergent alterations of spine densities and spine sizes.

### Whole-cell patch-clamp recordings

To probe the effects of NBQX administration at different concentrations on synaptic transmission, whole-cell patch-clamp recordings were conducted in CA1 pyramidal neurons. Recordings were performed at 35*^◦^*C. The bath solution contained (in mM) 126 NaCl, 2.5 KCl, 26 NaHCO_3_, 1.25 NaH_2_PO_4_, 2 CaCl_2_, 2 MgCl_2_, and 10 glucose (aCSF) and was continuously oxygenated with carbogen (5% CO_2_/95% O_2_). Glass patch pipettes had a tip resistance of 4 *−* 6 MΩ, filled with the internal solution which contained (in mM) 126 K-gluconate, 10 HEPES, 4 KCl, 4 ATP-Mg, 0.3 GTP-Na_2_, 10 PO-Creatine, 0.3% (w/v) biocytin. The internal solution was adjusted to pH = 7.25 with KOH and reached 290 mOsm with sucrose). We patched 6 neurons per culture to record the spontaneous excitatory postsynaptic currents (sEPSCs) of CA1 pyramidal neurons in the voltage-clamp mode at a holding potential of *−*70 mV. Series resistance was monitored before and after each recording and the neuron data was excluded if the series resistance went above 30 MΩ. Each neuron was recorded for 2 min.

### NBQX treatment

NBQX (2,3-dioxo-6-nitro-7-sulfamoyl-benzo[f]quinoxaline) is a competitive antagonist of AMPA receptors.^115^ We chose two concentrations 200 nM and 50 *µ*M and delivered by bath treatment to achieve a partial or complete inhibition of AMPA receptor currents. Wild-type cultures were either recorded in normal ACSF or ACSF that contained two different NBQX concentrations. In time-lapse imaging experiments, Thy1-eGFP cultures were treated with 200 nM and 50 *µ*M by adding NBQX (Cat. No. 1044, Tocris Bioscience, Germany) in the incubation medium for three days. Whenever we changed the new medium, fresh NBQX was administrated accordingly.

### Tissue fixation and immunohistochemical staining

Recorded cultures were fixed and stained for *post hoc* inspection. Cultures were fixed by immersing into 4% (w/v) paraformaldehyde (PFA) in 1*×* phosphate-buffered saline (PBS, 0.1 M, pH = 7.38) for 1 h and transferred into 1*×* PBS for storage at 4 *^◦^*C after being washed in 1*×* PBS. Before staining, all fixed cultures were again washed three times with 1*×* PBS (3 *×* 10 min) to remove residual PFA. We incubated the cultures with Streptavidin 488 (1 : 1000, #S32354, Invitrogen, Thermo Fisher, USA) in 1*×* PBS with 10% (v/v) in normal goat serum and 0.05% (v/v) Triton X-100 at 4 *^◦^*C overnight. In the next morning, cultures were rinsed with 1*×* PBS (3 *×* 10 min) and incubated with DAPI (1 : 2000) in 1*×* PBS for 20 min. After another 4 washes with 1*×* PBS (4 *×* 10 min), we mounted the cultures on glass slides with DAKO anti-fading mounting medium (#S302380 *−* 2, Agilent) for confocal microscope imaging.

### Time-lapse imaging

To inspect whether neural spine densities and spine sizes were altered by the three-day administration of NBQX, we employed time-lapse imaging to follow the same apical dendritic segments of CA1 pyramidal neurons before and after treatment. Live cell imaging was performed at a Zeiss LSM800 microscope with 10*×* water-immersion (W N-Achroplan 10*×*/0.3 M27; 420947-9900-000, Carl Zeiss) and 63*×* water-immersion objectives (W Plan-Apochromat 63*×*/1,0 M27; 421480-9900-000, Carl Zeiss). Thy1-eGFP tissue cultures where clear CA1 pyramidal neurons could be identified were used in this experiment. Dendritic segments from the radiatum layer were imaged. We imaged individual dendritic segments prior to the NBQX treatment and again after the three-day treatment. During the imaging session, the membrane insert with 4 cultures was placed into a 35 mm petri dish filled with 5 ml incubation medium on a platform which was constantly maintained at 35 *^◦^*C. We used the same pre-warmed and pH-adjusted incubation medium for imaging procedures.

### Experimental data quantification

Spontaneous excitatory postsynaptic currents (sEPSCs) were analyzed using the automated event detection tool from the pClamp11 software package as previously described.^116^

*z*-stacked fluorescent images of Thy1-eGFP cultures were projected to create a 2*D* representation of individual dendritic segments. ImageJ plugin Spine Density Counter^117^ was used to count spine numbers and measure segment length, which estimates spine density. For the same dendritic segments imaged at different time points, special attention was paid to ensure the same starting and ending points were used. *Post hoc* visual inspection was applied to ensure the spine detection results were not strongly biased. Both raw spine density and normalized spine density by baseline were used in the analysis.

The same *z*-projected fluorescent images were used to track individual spines for spine size analysis. To eliminate the bias from drawing and automatic reconstruction, we drew circles manually around the spine to cut it from the dendrite; the spine size was estimated by measuring the signal intensity with an arbitrary unit of the drawn circle. The drawing and measurements were performed with FIJI ImageJ. Both the raw values and normalized values by baseline spine size were used in the analysis. Statistical methods were specified in the individual results section.

In the following analysis, we normalized the alteration of spine sizes by their baseline size before treatment and binned the data by their initial sizes. We first fetched the spine sizes of individual spines pre-and post-treatment to calculate the alteration (Δspine size). Then, we normalized the alteration amount by the corresponding baseline spine size. We binned the spines by their baseline initial sizes in a third step. By sorting the spine ID in each binned group, we grouped the spine size alterations (Δspine size) by their initial spine size. This analysis shows us how spines with different initial sizes update their size over a certain time course with and without NBQX treatment.

### Statistical analysis

Dunn’s multiple comparison test was applied for statistical analysis regarding the sEPSC events among the three groups. For spine density analysis, the Wilcoxon test was applied to compare the values of each segment before and after the three-day treatment. If not otherwise stated, “ns” means no significant, “*” means *p <* 0.05, “**” means *p <* 0.01, “***” means *p <* 0.001. For spine size analysis, we first applied the Wilcoxon test for individual spines. To account for data clustering within each segment, linear mixed model (LMM) with *spine_size timing* was applied to compare the values before and after treatment with the segment ID as group factor as used in.^48^ The significance of LLM results was judged by whether the confidence interval (CI) crossed zero.

### Neuron model and network model

We used the same spiking neuron model and network architecture as described before in^46^ and in.^19, 48^ Current-based leaky integrated-and-fire point neuron was used for both excitatory and inhibitory neurons. We build an inhibition-dominated network with 10 000 excitatory neurons and 2 500 inhibitory neurons.^118^ To simplify the scenario, we only grow the connections within the excitatory population (E-E) with the activity-based structural plasticity rule (see the Structural plasticity rule section below). Each inhibitory neuron was beforehand hard-wired randomly to receive synapses from 10% of the excitatory and inhibitory population. All details and parameters concerning neural and network models can be found in the Supplementary Materials. We performed the network simulations with NEST simulator 2.20.2 and NEST 3.0^119^ and MPI-based parallel computation. All the model parameters and protocols can be found in Supplementary Tables 1-7.

### Structural plasticity rule

We enabled the growth, retraction, and rewiring of synapses among excitatory neurons with the help of structural plasticity rules. By definition, each excitatory neuron has multiple dendritic spines and axonal boutons, which are called synaptic elements. Synapses were formed by randomly matching free compatible synaptic elements. The growth and retraction of synaptic elements, or in other words, the number of synaptic elements, is governed by a growth rule. Three structural plasticity rules were explored in the current study: (i) linear growth rule; (ii) Gaussian growth rule with a zero setpoint and a non-zero setpoint; (iii) Gaussian growth rule with two non-zero setpoints. All three rules are determined by a function of calcium concentration that reflects neural activity,

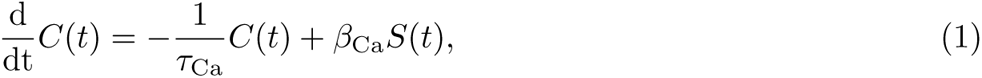

where *C*(*t*) is the time evolution of calcium concentration. Calcium concentration decays with a time constant *τ*_Ca_ and increases with calcium influx (*β*_Ca_) upon the emission of an action potential *S*(*t*) of the postsynaptic neuron. This operation is performed internally by the NEST simulator and works as a low-pass filtered signal of the spiking activity of the neuron. The growth of synaptic elements is regulated differently depending on the calcium concentration and the shape of the growth rule.

### Linear growth rule

The linear rule was first introduced in^45^ and systematically studied in an inhibitory-dominant neural net-work.^19, 46–48^ The number of synaptic elements (*z*(*t*)) is linearly dependent on the calcium concentration,

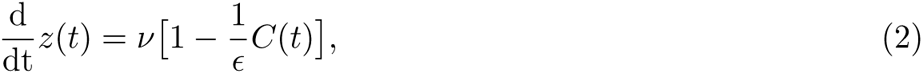

where *ν* is the growth rate, and *ɛ* is the target level of calcium concentration. Since calcium concentration reflects the neural activity loyally, the target level also suggests a setpoint of firing rate in the context of a certain neural network. As discussed before, when neurons fire below their target rate, they grow new synaptic elements and form new synapses. On the other hand, they break existing synapses and retract synaptic elements when they fire above the target rate (setpoint).

### Gaussian growth rule

The Gaussian rule has a more complex dependency on the calcium concentration when neural activity is too low, as shown in Equation 3. This rule was suggested and explored initially in.^43^

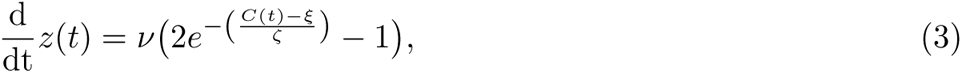

where 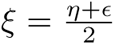 and 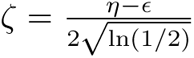. In this rule, *η* and *ɛ* are two setpoints: *ɛ* is the stable setpoint as used in the linear rule. When the neuron fires above *ɛ*, it retracts synaptic elements as in the linear rule. *η* is another setpoint introduced specifically for the Gaussian rule, determining the regulation manner when the neuron activity drops below *ɛ*. When the neuron is firing below *ɛ* but above *η*, the number of synaptic elements present will undergo homeostatic outgrowth, but when the neuron is firing below *η*, neurons will break synapses and retract elements. In the case where *η* = 0, neurons cease to change synaptic elements when their firing rate drops to zero.

### Homeostatic synaptic scaling

In order to achieve homeostatic synaptic scaling, we make use of a new synaptic model in NEST called *scaling_synapse*. In this synapse model, the weight of the synapse is regulated by the difference between a homeostatic setpoint and the calcium trace of the postsynaptic neuron,

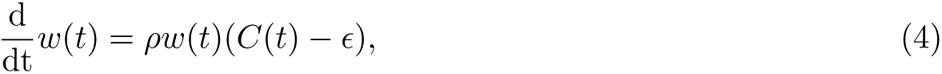

where *ρ* is the scaling factor, and *ɛ* is the same target value as used in the homeostatic structural plasticity rule.

### Activity perturbation

To examine different activity-induced scenarios on neural network connectivity, we performed systematic activity manipulation to a subnetwork of excitatory neurons (*N*_sub_ = 1 000), by changing its Poissonian input from 0% to 200% fold of the original intensity (FOI). All manipulations were performed at 6 000 s when the network has grown to the equilibrium state respectively with three structural plasticity rules. For the Gaussian rule with two non-zero setpoints, we applied damping current injection to the soma to facilitate its growth within the first 4 000 s and the network was simulated for another 2 000 s without any facilitating current.

### Quantifying firing rate, network connectivity, and synapse number

#### Firing rate

Firing rate was calculated by the average spike count over a recording period for individual neurons. We have long interval (5 s) and short interval (1 s). Short intervals were only used within the short time window after activity perturbation to reveal its transient dynamics; otherwise, long intervals were used.

#### Network connectivity

Two types of connectivity were used in the present study. We used a *N × N* connectivity matrix (*A_ij_*) to represent the recurrent excitatory connections of our network, where columns and rows correspond to pre-and postsynaptic neurons. For structural connectivity, the entry *A_ij_* of the matrix represents the total number of synaptic connections from neuron *j* to neuron *i*. For effective connectivity, we integrated the synapse number with individual weights for each pair of neurons by summating the total weights. So the entry of the connectivity matrix is the equivalent number of unit synapses, by dividing the sum with a uniform weight 0.1 mV. To average the mean connectivity of the whole network or a subnetwork at any given time *t*, corresponding columns and rows of the connectivity matrix were selected and averaged by 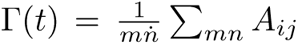. Synapse numbers were calculated by the sum of the entry in the structural connectivity matrix.

#### Synapse number

Input and output synapse numbers, also called indegree and outdegree in another context, were calculated by summating the input and output synapse numbers of individual excitatory neurons based on the structural connectivity matrix.

#### Quantifying the discrepancies in firing rate and connectivity from the target values

To apply a systematic comparison among different combinations of the synaptic scaling strengths and structural plasticity growth rates, we summated the discrepancies in firing rate and connectivity from the target values for the subpopulation over time as an index of activity and connectivity recovery. All the discrepancies were calculated by estimating the area between the actual time course and the equilibrium connection probability (10%) or the target rate (*ɛ*) from the time of silencing until the end of the simulation. The method was explained in detail in Supplementary Fig. 6. To achieve the heat maps in Fig. 9, we averaged 11 random trials for each parameter combination.

## Supportive Information

The authors declare that they have no conflict of interest.

## Acknowledgments

The work was supported by Deutsche Forschungsgemeinschaft (DFG; Project-ID 259373024 B14–CRC/TRR 167 to AV). The research leading to these results has received funding from the European Union’s Horizon 2020 Framework Programme for Research and Innovation under the Specific Grant Agreement No. 945539 (Human Brain Project SGA3). This research has also been partially funded by the Helmholtz Association through the Helmholtz Portfolio Theme Supercomputing and Modeling for the Human Brain. We acknowledge the use of Fenix Infrastructure resources, which are partially funded by the European Union’s Horizon 2020 research and innovation program through the ICEI project under grant agreement No. 800858.

## Author contributions

H.L. and A.V. conceived the project. M.L., H.L., and A.V. designed the experiments. M.L. performed the whole-cell patch-clamp recordings and analyzed the recorded data. H.L. performed the time-lapse imaging experiments and analyzed the spine data. H.L. established the bi-phasic HSP model in the network simulation and performed systematic explorations. S.D. established the HSS model and integrated it into the HSP model. H.L. and S.D. performed numerical simulations for the interaction between the two models. H.L. made the figures and drafted the manuscript. All authors contributed to the revision.

## Additional information

**Accession codes**: All raw data, simulation code, and analysis scripts are available here https://github.com/ErbB4/HSP-SS-interplay.

## Notes

https://github.com/ErbB4/HSP-SS-interplay

### Competing Interest Statement

The authors have declared no competing interest.

### Summary of Updates

We reorganized our text and figures to compare computer simulation results side-by-side with experimental results and spent tremendous core hours repeating another ten random trials of simulations included in the parameter scan experiment to achieve an averaged smooth output.

