## supplementary materials for "The interplay between homeostatic synaptic scaling and homeostatic structural plasticity maintains the robust firing rate of neural networks"

**Supplementary Table 1:** Parameters for the point neuron model

| $\tau_m$ | $t_{\mathrm{ref}}$ | $V_0$ | $V_{\mathrm{reset}}$ | $V_{\mathrm{th}}$ | $C_{\mathrm{mem}}$ |
|---|---|---|---|---|---|
| 20.0 ms | 2.0 ms | 0.0 mV | 10.0 mV | 20.0 mV | 250 pF |

**Supplementary Table 2:** Parameters for the network model

| $N_{\mathrm{E}}$ | $N_{\mathrm{I}}$ | $\Gamma_{\mathrm{E-I}}$ | $\Gamma_{\mathrm{I-E}}$ | $\Gamma_{\mathrm{I-I}}$ | $J_{\mathrm{E}}$ | $J_{\mathrm{I}}$ | $r_{\mathrm{ext}}$ |
|---|---|---|---|---|---|---|---|
| 10 000 | 2 500 | 10% | 10% | 10% | 0.1 mV | −0.8 mV | 30 kHz |

Varied values for $r_{\mathrm{ext}}$ were used for activity perturbation.

**Supplementary Table 3:** Parameters for the linear structural plasticity model

| $\epsilon$ | $\nu$ | $\tau_{\mathrm{Ca}}$ | $\beta_{\mathrm{Ca}}$ |
|---|---|---|---|
| 0.0079 | 0.00395 $\mathrm{s}^{-1}$ | 10 s | 0.0001 |

**Supplementary Table 4:** Parameters for the stable Gaussian structural plasticity model

| $\epsilon$ | $\eta$ | $\nu$ | $\tau_{\mathrm{Ca}}$ | $\beta_{\mathrm{Ca}}$ |
|---|---|---|---|---|
| 0.0079 | 0 | 0.004 $\mathrm{s}^{-1}$ | 10 s | 0.0001 |

**Supplementary Table 5:** Parameters for the bi-phasic Gaussian structural plasticity model

| $\epsilon$ | $\eta$ | $\nu$ | $\tau_{\mathrm{Ca}}$ | $\beta_{\mathrm{Ca}}$ |
|---|---|---|---|---|
| 0.0079 | 0.0007 | 0.004 $\mathrm{s}^{-1}$ | 10 s | 0.0001 |

**Supplementary Table 6:** Parameters for the synaptic scaling model

| $\epsilon$ | $\rho$ |
|---|---|
| 0.0079 | [0,0.1] |

**Supplementary Table 7:** Protocols for numeric stimulation

| | plasticity rules | $\eta$ | growth rate | $I_{\text{facilitating}}$ | FOI | $\tau_{\text{Ca}}$ | $\rho$ |
|---|---|---|---|---|---|---|---|
| Fig. 3Bi,Di | linear | - | 100%, 50%, 10% | | | 10 s | - |
| Fig. 3Bii,Dii | Gaussian | 0 | 100%, 50%, 10% | | | 10 s | - |
| Fig. 3Biii,Diii | Gaussian | 0.0007 | 100%, 50%, 10% | | | 10 s | - |
| Fig. 4A-C | Gaussian | 0.0007 | 100% | | | 10 s | - |
| Fig. 4E | Gaussian | 0.0007 | 100% | 750 pA | | 10 s | - |
| Fig. 4F | Gaussian | 0.0007 | 100% | 0, 200, 500, 750 pA | | 10 s | - |
| Fig. 5B,C | Gaussian | 0.0007 | 100% | 750 pA | 110%, 95%, 0% | 10 s | - |
| Fig. 5D | Gaussian | 0.0007 | 100% | 750 pA | [0%, 200%] | 10 s | - |
| Fig. 5E | - | - | - | - | - | 1 s, 10 s | - |
| Fig. 5F, orange | Gaussian | 0.0007 | 100% | 750 pA | 0% | 1 s | - |
| Fig. 5F, dark blue | Gaussian | 0.0007 | 100% | 750 pA | 0% | 10 s | - |
| Fig. 5F, light blue | Gaussian | 0.0007 | 10% | 750 pA | 0% | 10 s | - |
| Fig. 6B,C | Gaussian | 0.0007 for D, 0.0004 or 0.001 for A | 10% | 750 pA | 0% | 10 s | - |
| Fig. 6E,F | Gaussian | 0.0007 | 10%, 100% | 750 pA | 0% to 100% or 0% to 200% | 10 s | - |
| Fig. 7B-F | Gaussian + scaling | 0.0007 | 10% | 750 pA | 0% | 10 s | 0, 0.01, 0.02 |
| Fig. 8C-F | Gaussian + scaling | 0.0007 | [0%, 100%] | 750 pA | 0% | 10 s | [0, 0.1] |

The column **growth rate** summarizes the percentages of the original growth rate $\nu$ used for structural plasticity.

The column **FOI** summarizes the percentages of original intensities used for external activity perturbation.

For Fig. 5D, FOI values ranging from 0% to 200% was used for the systematic study.

Fig. 5E was performed in a single neuron model without any plasticity.

In Fig. 6BC, distinct $\eta$ values were used for axonal and dendritic elements. Otherwise, the same value was used for both parts.

In Fig. 6E,F, silencing protocol was applied (0% FOI) and then external stimulation was applied with 100% or 200% FOI.

In Fig. 7B-F, only three values for scaling factor ($\rho$) were used.

In Fig. 8C-F, a series values for $\rho$ ranging from 0 to 0.1 100%$\nu$ and a series values for growth rate ranging from 0%$\nu$ to 100%$\nu$ were used for the systematic study.

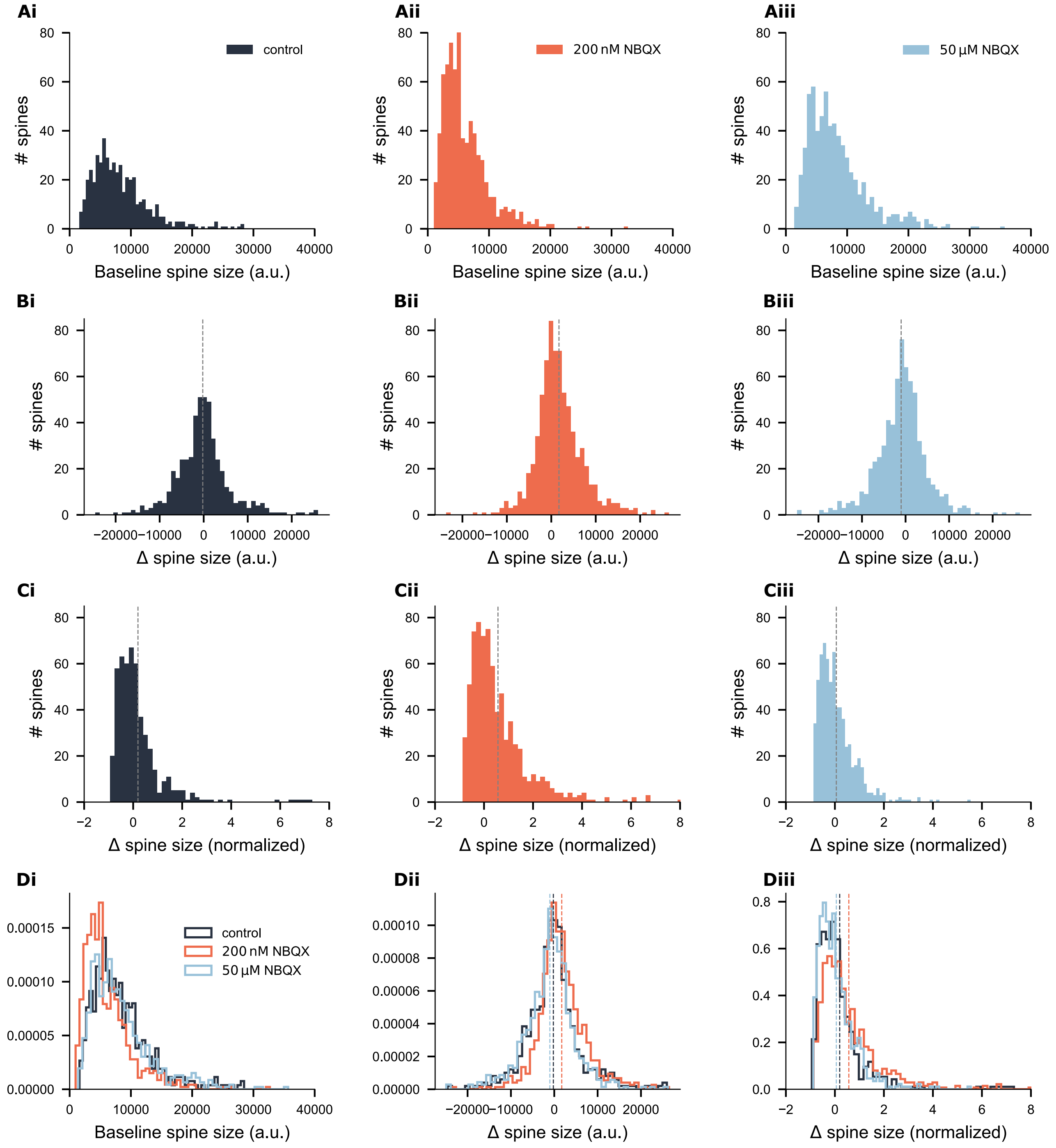


**Supplementary Fig. 1: Distributions of baseline spine sizes and changes of spine sizes over a three-day course in all three groups.** (A) Distributions of baseline spines sizes before treatment. (B-C) Distributions of raw size changes and normalized changes by the initial spine sizes. Vertical lines indicate the corresponding mean values. (D) Probability density functions that summarize the corresponding values in all three groups. Vertical lines with corresponding color codes indicate the mean values for each group.

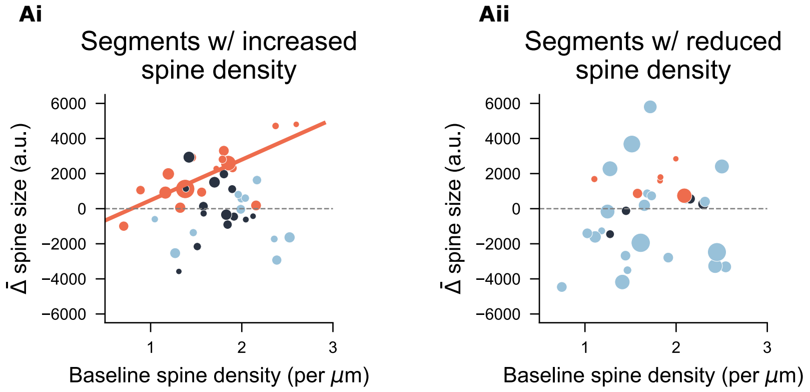


**Supplementary Fig. 2: Each segment's average change in spine sizes against its initial spine density.** This figure splits the data presented in Figure 2G by separating segments into two categories: spine density was increased or decreased. The marker size labels the net change in spine density over a three-day course. A positive correlation was observed in the 200 nM NBQX-treated segments with increased density ($r(18) = 0.74$, $p = 0.0005$, Pearson's correlation).

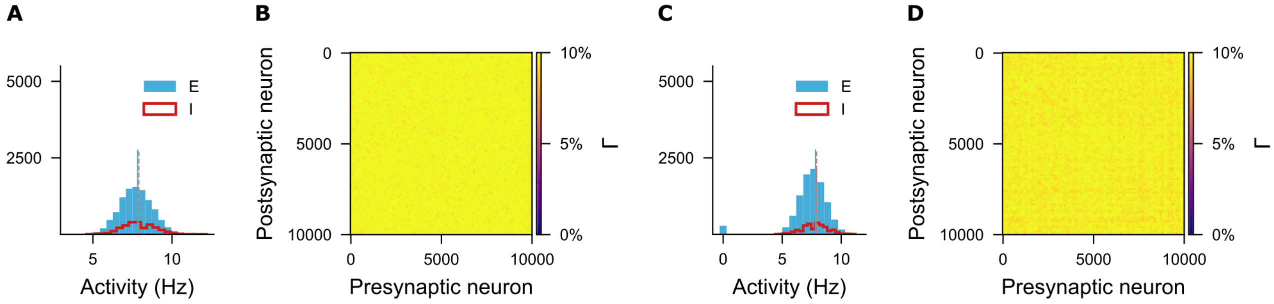


**Supplementary Fig. 3: Firing rate distributions and connectivity matrices at the indicated time points in Fig. 3.** A and B are for the linear growth rule. C and D are for the stable Gaussian rule ($\eta = 0$).

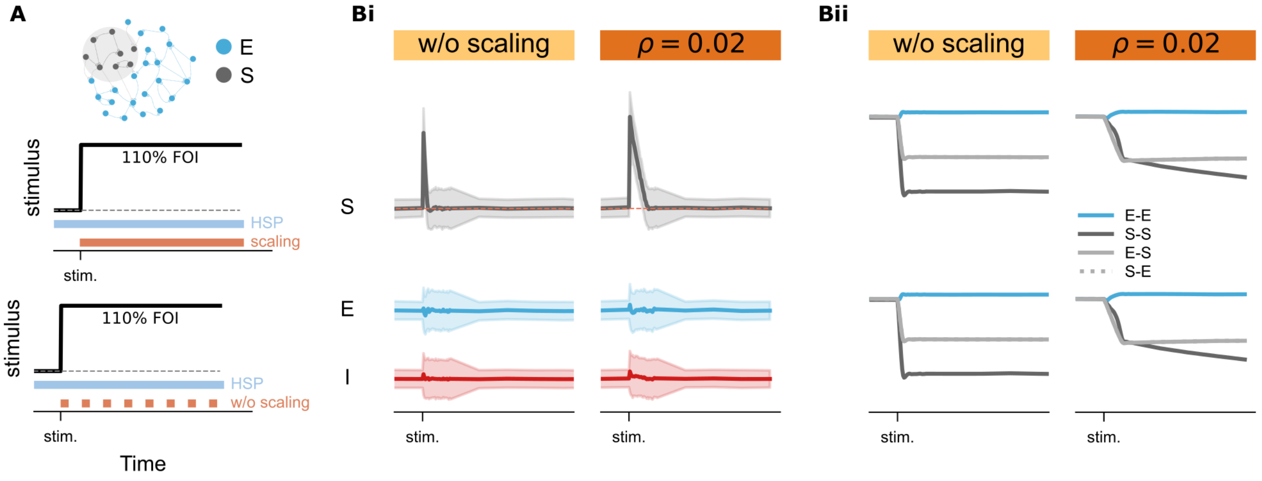


**Supplementary Fig. 4: Time courses of network activity and structural and effective connectivity under stimulation protocol with and without synaptic scaling rule.** $\rho = 0.02$ was the scaling factor used here and 110% FOI was used for stimulation.

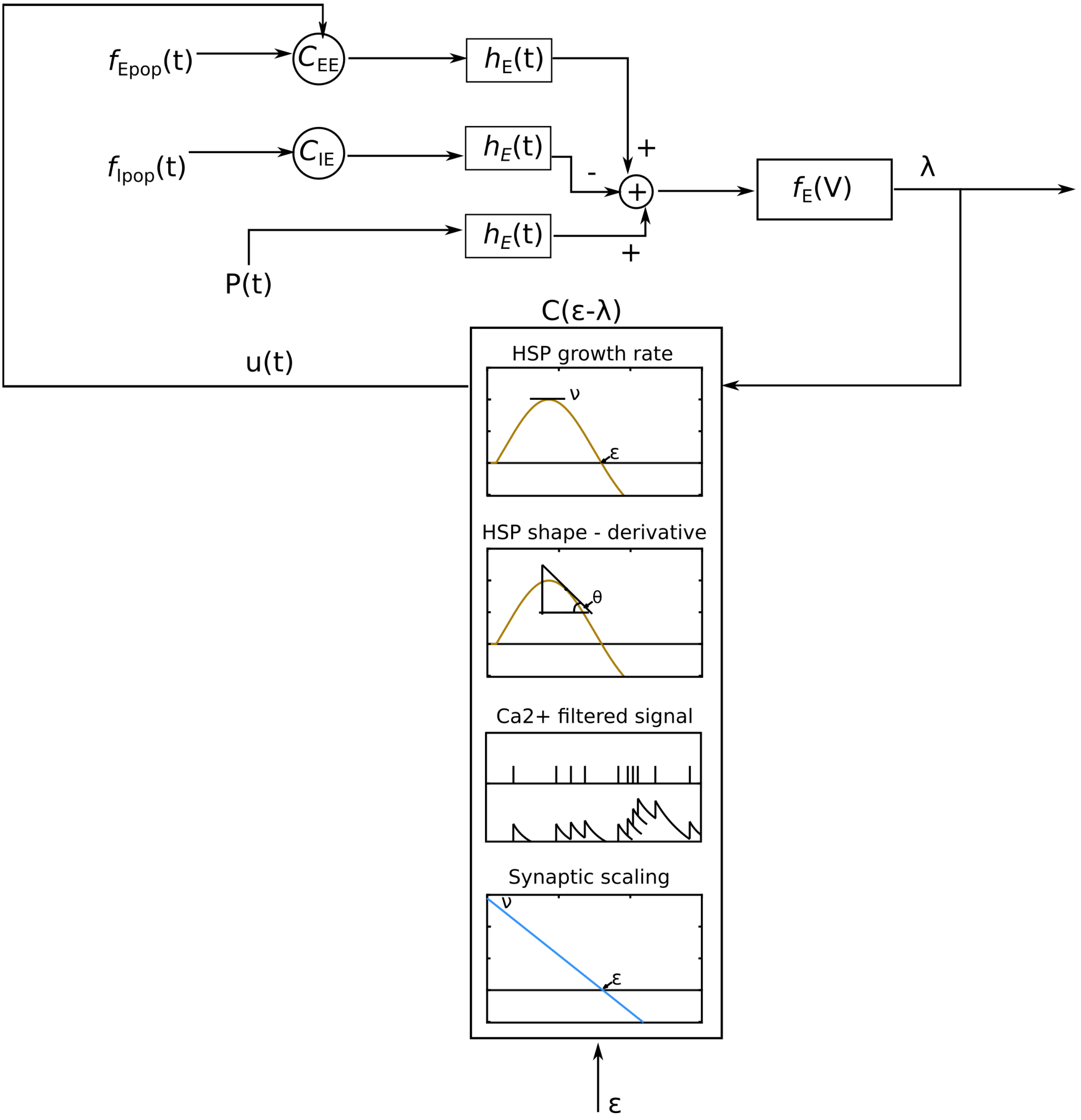


**Supplementary Fig. 5: Diagram describing the plasticity rules used in this manuscript in terms of control theory.**

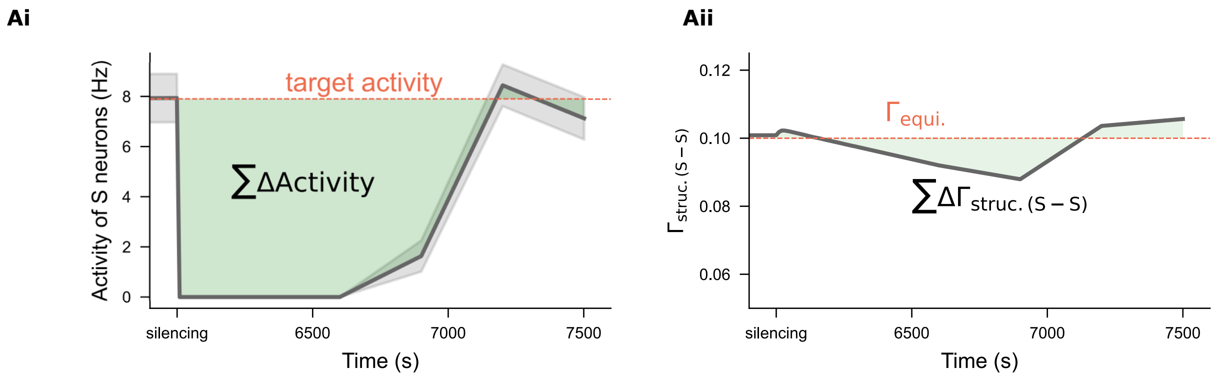


**Supplementary Fig. 6: Quantifying** $\Delta$Activity of S neurons **and** $\Delta\Gamma_{\text{struc.(S-S)}}$ **for plotting the heatmap in Fig. 8E,F.** Orange lines label the target values for firing rate and structural connectivity. The discrepancies in firing rate and connectivity were calculated and summated up the area over several time steps after switching on the silencing protocol.
